# Isolation and Characterization of Strains used in Bacterial-Based Strategies for Accelerated Carbonation of Lime Mortars

**DOI:** 10.1101/2025.04.10.648092

**Authors:** Franco Grosso Giordano, Quinten Mariën, Nele De Belie, Carlos Rodriguez-Navarro, Nico Boon

**Author notes:** **Email addresses**.

## Abstract

Over the last century, lime has been quickly replaced by the uptake of Portland cement, mainly due to its faster hardening. Achieving earlier hardening in lime through faster carbonation is thus essential to help overcome one of lime’s limiting qualities. In this work, we isolated and selected strains suitable for use in lime mortars, and used bacterial suspensions to carbonate lime materials. An isolation campaign from a lime mortar wall returned two alkaliphilic isolates, *Shouchella clausii* and *Shouchella patagoniensis. S. clausii* was then further adapted to high pH (> 11) by adaptive laboratory evolution to produce a third strain. All three strains were then followed for a period of 14 days in serum bottles at pH 11 and gas composition of the headspace, intact/damaged cell populations and pH were measured. In parallel, lime mortar samples were incubated in a closed environment with bacterial suspension of the strains. The mortars were then tested at 7 and 14 days with thermogravimetric analysis to study the amount of carbonation through bacterial activity. Overall, *S. patagoniensis* produced more CO_2_, close to the estimated maximum CO_2_ uptake rate of lime, and carbonated the lime mortars faster and to a larger extent than the other strains. Finally, the bacterial suspensions were directly mixed with lime and the carbonation front was followed using analytical techniques. Once again *S. patagoniensis* led to faster carbonation. Overall, it was shown that bacterial-based strategies for accelerated carbonation of lime are a possibility despite the high alkalinity of the binder.

**Importance:** Portland cement is the dominant binder used in most construction today, but until last century, lime was the ubiquitous construction material. The increase in use of cement has sprung from its higher strength and faster hardening, yet, lime still remain a relevant material, particularly in masonry structures and the built heritage. As such, novel lime materials are necessary to tackle some of the current limitations of lime, such as earlier hardening, which would not only make lime easier to work with but would also limit failure due to environmental conditions. As existing strategies to speed-up lime hardening have had limited uptake due to their reliance on expensive and often toxic chemicals, the need for novel solutions is in place. We show that bacterial-based strategies can help achieve a bio-based solution to go beyond the limitations of current strategies and open up new possibilities for microbial bioengineering in construction materials.

## Introduction

The last century saw a rise in the use of Portland cement as the dominant construction material, and with it, its use in built heritage restoration (1). Yet, the use of cement for the conservation and repair of these historical structures poses a huge risk as the material is not compatible with the masonry structures. For example, the higher porosity of lime mortars as compared to cement mortars allows for better conduction of moisture resulting in lower moisture or salt damage to masonry (2). Lime, a material used for millennia in masonry, is therefore more appropriate, yet it has some drawbacks. One of the main reasons for the increased adoption of cement instead of lime was due to the slow hardening process of lime (1).

The process of setting and hardening of aerial lime mortars is through the reaction of hydrated lime with

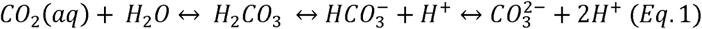

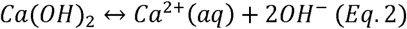

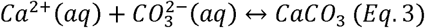

The main rate limiting step in the overall carbonation of hydrated lime mortars is the diffusivity of CO_2_ to the pore system where the dissolution of CO_2_ will occur (Eq. 1) (3). In atmospheric conditions, this reaction can be very slow with reports of 1700 year old mortars that are still not fully carbonated (4). CO_2_ diffusivity can be affected by the pore size distribution of the matrix and the water content. Small pore size leads to lower gas exchange whereas a high water saturation directly inhibits the diffusion of CO_2_ as its diffusion rate can be 10,000 times lower than in air (5). Moreover, as carbonation occurs from outside to inside the material, the deepest sections of the matrix take longer to carbonate (3). Under ideal conditions, i.e. a thin layer of lime putty paste exposed to high CO_2_ concentrations as tested by Van Balen (5), CO_2_ uptake per unit time reached an average maximum ranging from 3.99 x 10^-4^ to 2.84 x 10^-3^ mmol of CO_2_ absorbed per mmol of lime per second (1/s). It follows that one strategy to overcome the slow hardening of lime-based mortars would involve speeding up carbonation to such rates. Hence, shifting the equilibrium of Eq. 1 to the right is essential to speed up carbonation at an early age.

Three parameters can be influenced to improve carbonation rates: increasing the temperature, limiting humidity to an optimal range of 40 to 80 % (6) and increasing CO_2_ concentration ([CO_2_]) up to approximately 20 % (7). Of all three, the latter is the most studied and strategies have been engineered to use in lime materials. It can be affected through increasing diffusivity of CO_2_, by increasing locally the partial pressure of CO_2_ (*p*CO_2_) and/or CO_2_ concentration in air, or increasing the dissolution of CO_2_ in water to produce bicarbonate ions (Eq. 1). Increasing CO_2_ diffusivity can be achieved by changing the microstructure of the material to allow faster airflow (i.e., by increasing the porosity, pore size, and pore connectivity). Nevertheless, this can lead to detrimental changes in mechanical properties and the outside-to-inside process of carbonation remains a hurdle. Changes in *p*CO_2_ or CO_2_ concentration, usually achieved in controlled environments, are effective up to a maximum, ranging from 5 % (8) to 20 % [CO_2_] (7), after which carbonation can lead to the formation of an impervious layer of CaCO_3_ that completely inhibits the progress of carbonation (5). Also, changes in *p*CO_2_ are heavily impacted by humidity (9), as the dissolution/hydration of CO_2_ (Eq. 1) is the governing rate limiting step in the lime carbonation. Lastly, too high a concentration of CO_2_ can lead to a high heat release from the exothermic reaction of carbonation leading to premature drying and the stopping of carbonation (10). All this highlights that optimal solutions for carbonation need to strike a balance between the increase in CO_2_ availability through the entire matrix while maximizing the dissolution of CO_2_.

Multiple strategies for enhancing carbonation already exist. Of the many strategies implemented (3), the most positive results in terms of accelerating carbonation have been observed with some additives, especially organic ones, such as vegetable matter (11), plant extracts (12) and animal products (13). A little explored strategy is, however, the use of microbial-mediated carbonation. In principle, microorganisms possess multiple qualities that make them attractive as early hardening additives. Firstly there is the use of microbially-induced carbonate precipitation (MICP). In MICP, metabolites from microbial activity – of which those primarily studied have been bacterial – generally lead to the formation of CO_3_ ^2-^ ions which can then react with free Ca^2+^ to form CaCO_3_ (Eq. 3) (14). One of these pathways is the release of CO_2_ through the breakdown of organic acids during respiration. This can be used to increase the rate of carbonation with the possibility of creating a CO_2_ source working from the inside of the material. Lopez-Arce et al., showed that such a strategy was also possible using a yeast fermentation system - another microbial source of CO_2_ different to bacteria - to carbonate Ca(OH)_2_ nanoparticles applied for the consolidation of limestone samples placed in a separate container (15). Moreover, a patent application exploring microbially-mediated carbonation saw positive results when directly mixing a bacterial suspension with the mortar, achieving at least 1.2 times higher compressive strengths than reference mortars (16). Still, here the addition of urea or organic acids was the biggest enhancer of the compressive strength, while bacteria activity led to a small cumulative increase. Secondly, another benefit is that the CO_2_ release by microorganisms is not instantaneous, preventing the immediate precipitation of CaCO_3._ This is a problem in CO_2_ sources derived from chemical compounds, as it can be detrimental to the fresh properties (rheology) of lime materials. For example, acceleration of carbonation can be achieved using carbamates which in contact with water decompose releasing CO_2_ right upon mixing with the fresh lime paste (17). The study of carbamates has nonetheless recently been revived and could be promising in the future (18). Thirdly, the diversity of microbes, in particular bacterial species, offers a large array of options to optimize microbially-mediated carbonation, through the choice of metabolic pathways and selection of microbial growth kinetics (14). Finally, bacteria also release extracellular polymeric substances (EPS), composed of a mixture of proteins, polysaccharides and other organic compounds. EPS have been observed to control the morphology and kinetics of the biomineralization of CaCO_3_, and this has further effects on the structure-function of calcite (19). Once again, this further suggests that microbial-mediated carbonation of lime is a sensible strategy.

It is worth noting that bacterial additives have been previously engineered for construction materials. For example, work on bio-additives in cement mixtures is very well established for bioconsolidation or as self-healing agents (20). Similar hurdles are faced in terms of application in fresh cement as in fresh lime, particularly in terms of incorporation of the bacterial additives to the cement mixtures. The cement environment can be very harsh due to pH reaching >12 in a fresh state as well as the small pore size of the cement matrix. For this reason, protective mechanisms for the bacteria are often implemented which improves survivability in the long term while alkaliphiles are more often selected to better survive in the high pH environment (21). MICP has also been used for consolidation of limestone to restore and improve lost mechanical properties due to weathering through the creation of new calcium carbonate cement by applying bacteria (22, 23) or through the activation of the indigenous carbonatogenic bacteria already present in the stone substrate (24). Nevertheless, the conditions of application in these works have been very different from those expected in fresh lime mortars, specifically due to the close-to-neutral pH of limestone and the superficial application of bacteria.

As such, inspired in the engineered process of MICP used in cement and concrete, here we study the potential of bacterial-based strategies for early age hardening of hydrated lime. The approach taken seeks to increase the availability of CO_2_ through the bacterial metabolism of organics, which should speed up Eq. 1, leading to a faster carbonation in the presence of bacteria than without them. This work serves as a proof-of-concept for such a strategy. Importantly, the focus is doing so at an early age (up to 14 days), which would produce faster setting lime mortars, overcoming the main limiting quality for the wider use of hydrated lime mortars (3). Therefore, we performed an isolation campaign from which two novel strains were selected and studied in-depth for their alkalophilicity, their CO_2_ production rates, their capacity to carbonate lime mortars and lastly, their effect on hydrated lime when directly incorporated into the material.

## Materials and methods

### Environmental sample

A lime mortar from a wall in Ghent, Belgium was sampled (Figure 1). A part of the mortar sample was ground and analysed under X-ray diffraction (XRD) to disclose its mineralogy with a Siemens D5000 (Munich, Germany) (measurement parameters: Cu-Kα radiation, 40 kV, 40 mA, 5 to 90 °2θ exploration range). After analysis, the mortar was placed in a 37 % HCl solution and stirred for 30 min. The suspension was filtered, and the acid insoluble residue was collected and washed thrice. XRD was performed again on the residue. The XRD analysis confirmed the mortar was indeed a lime mortar, composed of calcite (as binder) and quartz (as aggregate) (Figure S 1).

**Figure 1.**
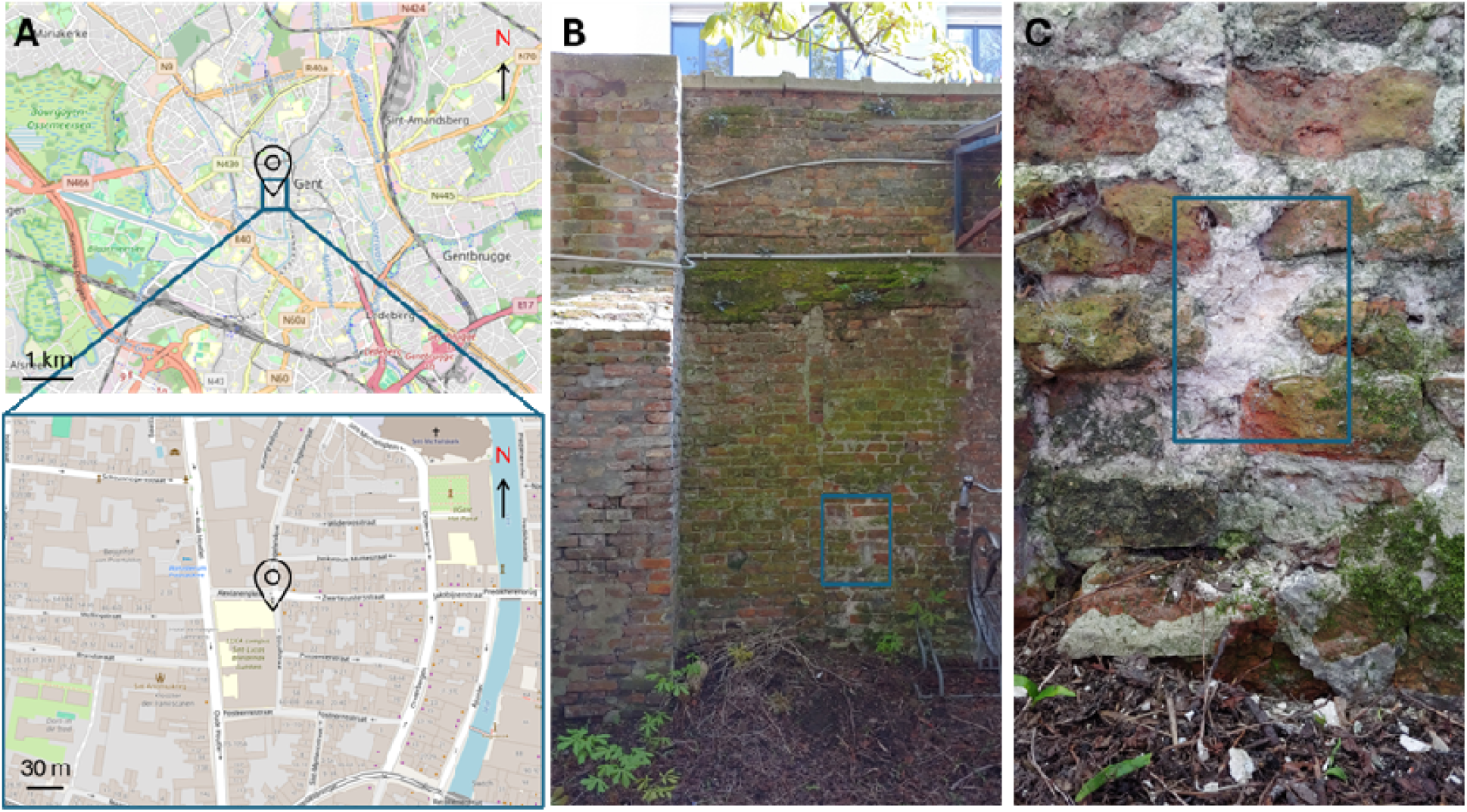
A) Top: Location of sampling of lime mortar masonry wall in Gent, Belgium. Bottom: Zoom-in of location o the eastern wall. B) Front view of the sampling area (blue square) on the north-east wall, where the least amount of biological degradation was observed. C) Zoom-in of the cleaned wall prior to sampling the internal lime mortar (blue square).

The mortar was then placed under UV light for 10 minutes on all sides to sterilize the surface. Using a surface sterilized mortar and pestle the lime mortar was ground and media were inoculated with mortar. A neutral (pH 7.0 – 7.4) low nutrient medium, R2A (Yeast Extract 0.5 g/L; Proteose Peptone 0.5 g/L; Casein Hydrolysate 0.5 g/L; Glucose 0.5 g/L; Soluble Starch 0.5 g/L; Na-pyruvate 0.3 g/L; K_2_HPO_4_ 0.3 g/L; MgSO_4_ anhydrous 0.024 g/L), and two high pH media, Horikoshi (Glucose 10.0 g/L, Polypepetone 5.0 g/L, Yeast Extract 5.0 g/L, K_2_HPO_4_ 1.0 g/L, MgSO_4_ x 7 H_2_O 0.2 g/L, 10 g/L Na_2_CO_3_; final pH 10) and R2A adjusted to pH 9.2 with carbonate buffer, were used. The enrichment was grown for 48 h and then diluted 10^-3^ to 10^-5^ and plated on agar plates of the same media. Morphologically distinct colonies were transferred to a new plate and streak cleaned 3 times to ensure purity of the strain. All samples were incubated at 28 °C between 24 h to 120 h.

### Microbe maintenance

All strains were grown and maintained in Tryptic Soy Broth (TSB: Pancreatic Digest of Casein 17.0 g/L, Peptic Digest of Soybean 3.0 g/L, Glucose 2.5 g/L, NaCl 5.0 g/L, di-Potassium hydrogen phosphate 2.5 g/L) or Agar (TSA: Pancreatic Digest of Casein 17.0 g/L, Peptic Digest of Soybean 3.0 g/L, NaCl 5.0 g/L, Agar 14 g/L). For each experiment, a double transfer was performed to reduce carryover effects from agar plates or stationary phases. All growth conditions were at 28 °C and 140 RPM unless stated otherwise.

### Isolate selection

Strains were first selected for their growth capacity at high pH. Isolates were inoculated to OD_600_ 0.01 in microtiter plates in TSB adjusted to pH 10 using carbonate buffer (TSB pH 10: Pancreatic Digest of Casein 17.0 g/L, Peptic Digest of Soybean 3.0 g/L, Glucose 2.5 g/L, NaCl 5.0 g/L, NaHCO_3_ 3.9 g/L, Na_2_CO_3_ 5.7 g/L). The best growing isolates were then grown in TSB adjusted to pH 11 using CAPS buffer (TSB pH 11: Pancreatic Digest of Casein 17.0 g/L, Peptic Digest of Soybean 3.0 g/L, Glucose 2.5 g/L, NaCl 5.0 g/L, CAPS 22.13 g/L) (TSB pH 11).

After reducing the number of isolates based on their alkalophilicity, the CO_2_ production rate of the best growing bacteria was measured in a serum bottle experiment in TSB. A 1:100 inoculation was made in fresh TSB, and the isolates were grown for 24 h to achieve full growth. The TSB suspension was centrifuged at 1500 RCF for 10 minutes and resuspended in fresh TSB. Serum bottles of 30 mL volume with 10 mL of the bacterial suspension were prepared, capped with butyl rubber stoppers and sealed. The samples were incubated under constant shaking. 1.5 mL of the headspace of the bottles were samples using a syringe and needle at 0, 7, and 14 days and CO_2_ levels were measured using gas chromatography (see below).

The selected isolates were then deposited at the Belgian Coordinated Collections of Microorganisms (BCCM) and the sequences were submitted to the Database Resources of the National Center for Biotechnology Information (25) and the accession GenBank Accession number provided.

### Adaptive laboratory evolution for high pH tolerance

Adaptive laboratory evolution of an isolate was performed on a microtiter plate. The selected strain was inoculated to OD_600_ 0.01 in microtiter plates containing 200 μL TSB pH 11, in triplicate. After growth the best growing replicate was reinoculated with a 1 % inoculum in 200 µL TSB pH 11. The experiment was carried out for 10 consecutive transfers. The adapted strain was deposited at the BCCM.

### Generation calculation based on OD_600_ measurements

To determine the number of generations (‘n’) in a bacterial culture based on OD_600_ measurements, we utilized the following equation:

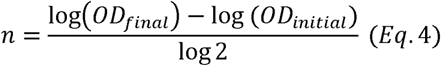

Where *OD_initial_*is the optical density at the beginning of the measurement period, *OD_final_*is the optical density at the end of the measurement period and *log2* accounts for the doubling nature of bacterial growth.

### Models and data analysis

All selected strains were grown in TSB pH 11 in 9 replicates each. The OD/time data was log-transformed and a Gompertz model was used for curve fitting for each growth curve. Outliers were removed based on Iglewicz and Hoaglin’s robust test for multiple outliers. The growth rate and the relative standard error (1σ) for each strain were estimated from the model for each strain.

### Metabolic kinetics and carbonation of lime mortar samples

The metabolic kinetics of the selected isolates were followed in a closed, glass serum bottle experiment (Figure S 2A). In parallel, and using the same bacterial suspension, the carbonation rate of lime mortar samples incubated in a close environment was followed (Figure S 2B). For the bacterial suspension, a 1:10 inoculation was made in fresh TSB pH 11. With the larger inoculum growth was achieved in 24 h to retain similar experimental time frames. The culture was centrifuged at 1500 g for 10 minutes and the pellet was resuspended in fresh TSB pH 11.

### Metabolic kinetics in a high pH environment

Four serum bottles were prepared with 10 mL of the bacterial suspensions resuspended in fresh medium, capped with butyl rubber stoppers and sealed. The samples were incubated and shaken. Sampling of 250 µL of bacterial suspension was performed from the serum bottle at 0, 1, 7, 10 and 14 days for pH measurement and flow cytometry analysis. 2 mL from the gas phase in the headspace of the bottles were sampled at 0, 1, 3, 5, 7, 10 and 14 days to measure O_2_ and CO_2_ levels. The headspace pressure was also measured with a tensiometer (GMH 3111 equipped with a 603310 MSD 2.5 BAE sensor, Greisinger) and needle. To test the aerobic requirements of the bacteria, 2 serum bottles were briefly reopened at day 7 of the experiment using a needle and filter, to ensure sterility, and the internal atmosphere of the bottle was stabilized with the external atmosphere. Serum bottles with only water and TSB pH 11 were prepared and handled equally as controls. One serum bottle also contained TSB + NaN_3_. NaN_3_ was used to inhibit bacterial growth for the experimental section on the carbonation of lime mortar samples (see below). Such control was also prepared here to follow the effect of NaN_3_ on the potential release of CO_2_ from the uninoculated media in the control lime mortar samples experiment. No effect was observed though (Figure S 3).

### Carbonation of lime mortar samples

A lime mortar mixture was prepared using a 1:3 lime:aggregate ratio (by volume) with a water:binder ratio of 0.85 (by volume). The aggregate used was a standard normalized quartz sand which fulfils the grain size distributions based on EN 196-1 (> 98 % SiO_2_). The hydrated lime was a CL 90-S (Lhoist, Belgium). The mortar was cast into a silicone mould containing wells of 1.4 x 1.4 x 1 cm^3^ (w x l x h). A layer of gauze and 2 layers of filter paper were placed on top and 1.5 kg of weight was applied to the mortar, adapting the standard mortar casting technique to smaller samples. They were left to rest for 1 h and then placed at 100 °C for 1.5 h for further drying.

After drying, 2 lime mortar samples were placed inside a Falcon tube separated by a layer of sterile aluminum paper from 9 g of sterilized cotton (Figure S 2B). Using a pipette, 10 mL of the bacterial suspensions resuspended in fresh medium were added to the cotton and then the tube was closed firmly. Tubes with cotton soaked with water (referred to as Water), tubes with cotton soaked with TSB pH 11 and 0.02 % NaN_3_ (referred to as TSB) and, tubes with a dry cotton (referred to as Air) were prepared as controls. At 7 and 14 days the soaked cotton was removed from the tube and the area was cleaned with a fresh cotton and ethanol, without coming into contact with the lime mortar samples. The soaked cotton was then replaced with clean, dry cotton and the lime mortar samples were further stored in the closed tubes. Two tubes per cotton suspension treatment and date (i.e., 4 per treatment) were prepared, All the tubes were stored inside a zip-lock bag. Soda lime with a CO_2_ indicator (Merch, Darmstadt, Germany) was placed in each tube to ensure proper storage and to halt any further mortar sample carbonation until thermogravimetric analysis.

### Flow cytometry

Total cell concentrations and intact/damage populations were measured as per (26). Briefly, a Attune™ NxT flow cytometer was used (ThermoFisher Scientific, Belgium) with BRxx configuration, equipped with a blue (488 nm, 50mW) and red laser (638 nm, 50mW), two fluorescence detectors with bandpass filters (BL1: 530/30 nm, BL3: 695/40 nm) and two scatter detectors on the 488 nm laser (FSC: 488/10 nm, SSC: 488/10 nm). The flow cytometer was operated with Attune™ focusing fluid (Thermofisher Scientific, Belgium) as sheath fluid. Samples were diluted between 10^-1^ and 10^-3^ prior to staining for intact/damaged population identification with a combination of SYBR® Green I (SG, 100 x concentrate in 0.22 µm-filtered dimethyl sulfoxide (DMSO), Invitrogen, Belgium) for intact cells and propidium iodide (20 mM in DMSO, Invitrogen, Carlsbad, CA) for damaged cells (SGPI). The samples were incubated for 20 minutes at 37°C in the dark. Samples were then analysed in triplicate in either fixed volume mode or maximum event count (20000 events) at a flow rate of 100 µL/min.

### Gas chromatography

The gas phase composition of the headspace was analysed with a Compact GC4.0 (Global Analyser Solutions, Breda, The Netherlands), with two channels. CO_2_ was measured on channel one, which was equipped with a Rt-QSBond precolumn and column. Channel one used an injection split flow of 2 mL/min, temperature of 70 °C and pressure of 110 kPa. Channel two was equipped with a Molsieve 5A pre-column and Porabond Q column (to measure O_2_ and N_2_). O_2_ and N_2_ were characterized using an injection split flow of 10 mL/min, temperature of 70 °C and pressure of 100 kPa. Helium was used as carrier gas for both channels and the column temperature was set to 50 °C. All gases were quantified with a thermal conductivity detector set to 110 °C with filament temperature of 210 °C and a reference flow of 100 kPa.

### Total CO_2_ production estimation

The CO_2_ concentration in the system or total inorganic carbon was estimated using the ideal gas law, Henry’s law, and the dissolution constants of CO_2_ (Eq. 5 - 12). The total inorganic carbon in the system (C_inorganic_ in mmol C/L) was estimated as the sum of CO_2_ in the headspace and all dissolved inorganic carbon species (i.e. [CO_2_]_aq_, H_2_CO_3_, HCO_3_ ^-^, CO_3_ ^2-^). The concentrations of dissolved species were derived from the headspace concentration assuming equilibrium between headspace and liquid according to the Henry constant, as well as equilibrium between all dissolved species according to the established dissociation constants (where *P* is the total absolute pressure in the system (atm), *p*CO_2_ is the partial pressure of CO_2_ (atm), m[CO_2_]_h_ is the measured concentration of CO_2_ in the headspace (%), [CO_2_]_h_ is the converted concentration of CO_2_ in the headspace (mol/L), [CO_2_]*_aq_*is the concentration of dissolved CO_2_ (mol/L), *T* is temperature (K), *V_h_* and *V_l_* is headspace and liquid volume (L), respectively, *n* is amount of substance (mol), *R* the ideal gas constant, *k* is Henry’s constant and, *K_H_, K_a1_,* and *K_a2_*, the dissolution constants of CO_2_):

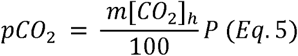

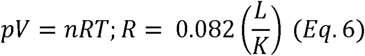

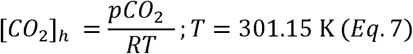

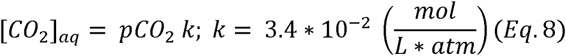

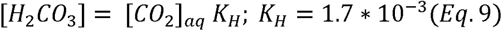

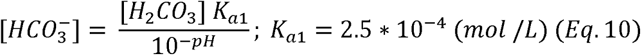

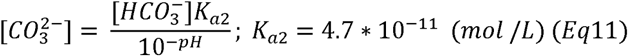

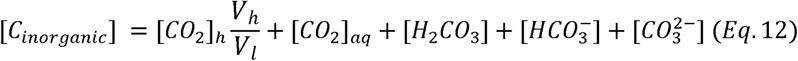

### Specific rate of O_2_ consumption

The O_2_ consumption per cell (mmol/L/cell), or specific rate of O_2_ consumption, allows for a better comparison between bacterial species in different initial conditions and provides information on cellular activity (27). It was calculated using the following formula:

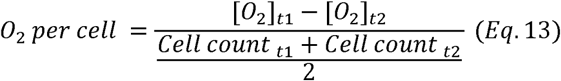

where the O_2_ concentration in the headspace ([O_2_] in mmol/L) between *t_1_*and *t_2_* is divided by the average total intact cell count between the two time points.

### Thermogravimetric analysis

Thermogravimetric analysis (TGA) was done in a TGA-1000-U (Navas Instruments, USA) for lime mortars. The whole sample (3.9 ± 0.2 g) was inserted into the machine and heated in a ceramic crucible in a N_2_ atmosphere, at 5 °C min^-1^, from 20 to 1000 °C. The mass loss was converted to percentage by normalizing to the initial total mass. To calculate the Ca(OH)_2_ content in the sample (CH), the percentage weight (wt %) loss between 350 and 550 °C (WL_CH_), molecular weights of Ca(OH)_2_ (74 g/mol) and water (18 g/mol) were used (Eq. 14 and 15). To determine the amount of CaCO_3_ in the sample (CC), the percentage weight loss between 650 and 950 °C (WL_CC_), molecular weights of CaCO_3_ (100 g/mol) and CO_2_ (44 g/mol) were used (Eq. 16 and 17). Finally, from these values, the percentage weight of the converted Ca(OH)_2_ (CH_conv_) was estimated, then the total initial Ca(OH)_2_ (CH_i_) was calculated and finally the carbonation percentage (Carb) (%) was calculated (Eq. 18 - 20):

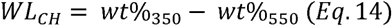

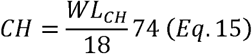

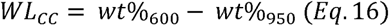

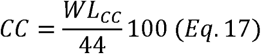

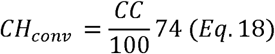

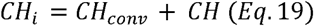

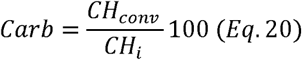

### Carbonation of lime pastes

To determine the effect of bacterial suspensions in lime, lime pastes were prepared. Lime pastes were then tested at 7, 12, 15, 21 and 35 days using TGA, Fourier-Transform Infrared Spectroscopy (FTIR) and phenolphthalein spraying.

### Preparation of lime pastes

The bacterial isolates were grown in TSB, and a 1:100 inoculation was made in fresh TSB. The isolates were grown for 24 h to achieve full growth. The TSB suspension was centrifuged at 1500 RCF for 10 minutes and resuspended in TSB without buffering agent (TSB-C: Pancreatic Digest of Casein 17.0 g/L, Peptic Digest of Soybean 3.0 g/L, Glucose 2.5 g/L, NaCl 5.0 g/L). Lime paste mixtures with either the resuspended bacteria or only TSB-C were prepared to study the capacity of the bacteria to carbonate the lime paste directly. Hydrated lime CL 90-S was used to prepare pastes using a 1:1 water-to-binder mass ratio, where the resuspended bacterial suspensions or only TSB-C were used in place of water. The paste was mixed by hand until homogeneous and cast into silicone moulds with wells of 1.4 x 1.4 x 1.0 cm^3^ (w x l x h). Due to the fragility of the pastes, 20 samples per mix were prepared to ensure that 3 samples were tested at each date. The moulds were tapped 30 times to remove internal bubbles and cured until the time of testing at 21 °C and 65 % RH. Leaving the samples in the mould meant that the carbonation front moved in a 1D, top-down direction.

### Thermogravimetric analysis

Three lime paste samples were grinded using a mortar and pestle. About 50 mg of powdered sample was then used to perform TGA in a Netzsch STA 449 Jupiter TGA-DTA Analyzer. The temperature ranged from 20 to 1100 °C at a heating rate of 10 °C/min in an inert nitrogen atmosphere. The carbonation percentage was calculated as above.

### FTIR spectroscopy

Fourier Transform Infrared Spectroscopy (FTIR) spectra of the precipitate were recorded on a NICOLET 20SXB FTIR (400–4000 cm–1 spectral range; resolution of 2 cm^−1^).

### Phenolphthalein analysis and measurement

A lime paste samples per mixture was split in half and sprayed with a 0.1 % solution of phenolphthalein pH indicator in ethanol. The indicator solution was allowed to develop for 1 h and images were taken of the samples through a microscope.

### Statistical analysis

Two-way analysis of variance (ANOVA) (α = 0.05) was performed to examine the effect of bacterial strains and time on the outcome variables (e.g., CO_2_ production, carbonation). Given significant differences in means (p < 0.05), we used Tukey’s Honest Significant Difference (HSD) test for post-hoc pairwise comparisons to determine which isolate groups differed. All tests performed in R.

## Results

### Isolation campaign

An isolation campaign was carried out to find strains with the goal of using them as carbonation enhancers in lime mortars. Given lime has a high pH and hardens through incorporation of CO_2_ the isolates should be fit to grow at high pH and produce high amounts of CO_2_ at an early age. To ensure they were adapted to the environment, a lime-mortar wall was chosen for the starting inoculum. A total of 33 isolates were obtained. Matrix-assisted laser desorption–ionization time-of-flight mass spectrometry (MALDI-TOF MS) (28) was used to dereplicate the strains through the comparison of spectral differences, which left 16 reference strains remaining (Table S 1). The most common genera were *Shouchella* (previously *Alkalihalobacillus*), *Bacillus* and *Staphylococcus*. Next, strains were excluded based on their classification as biosafety level 2, i.e. microbes that pose moderate hazards to humans and the environment, according to the BacDive database (29), leaving a final number of 12 strains.

### Strain selection

To find the strains most adapted to high pH environment, growth experiments were performed in TSB pH 10 and then at pH 11. Alongside the 12 isolates, *Lynsinobacillus fusiformis* (LMG 9816) was used as a control, as it has been commonly used in other studies of MICP and is known to be an alkaliphile (30, 31). Only five of the 12 selected isolates grew successfully at TSB pH 10 (Figure S 4A), with *L. fusiformis* failing to do so, and were tested for their capacity to grow in TSB pH 11. Four isolates grew at pH 11 (Figure S 4B) - isolates 2 and 3 reached maximum OD within 48 h, while isolates 14 and 13 did so at 60 h and 75 h, respectively.

All 4 isolates were studied further for their CO_2_ production capacity to be used as carbonation enhancers of lime mortars. Here a fully grown bacterial suspension in TSB was used as bacteria in the exponential phase will produce lower CO_2_ than those in the stationary phase (32). The fully grown suspension wa centrifuged and the pellet was resuspended in a fresh medium. The suspension was transferred to serum bottles and CO_2_ concentration was monitored over time. Isolates 2, 3 and 14 could be grouped as fast CO_2_ producers, showing a logarithmic production of CO_2_ that rose rapidly at an early age (Figure 2). Isolate 2 had a significantly higher CO_2_ production than isolate 3 (*p <* 0.05) but not than isolate 14. A such isolate 3 was excluded from further study. The isolates were sequenced, with isolate 2 being referred to as *Schouchella clausii* (LMG 33235, GenBank Accession No. PV163962) and isolate 14 a *Schouchella patagoniensis* (LMG 33400, GenBank Accession No. PV163981). These two isolates were the best adapted to high pH and were the fastest CO_2_ producers. They were therefore selected for a more in-depth characterization in a more realistic environment.

**Figure 2.**
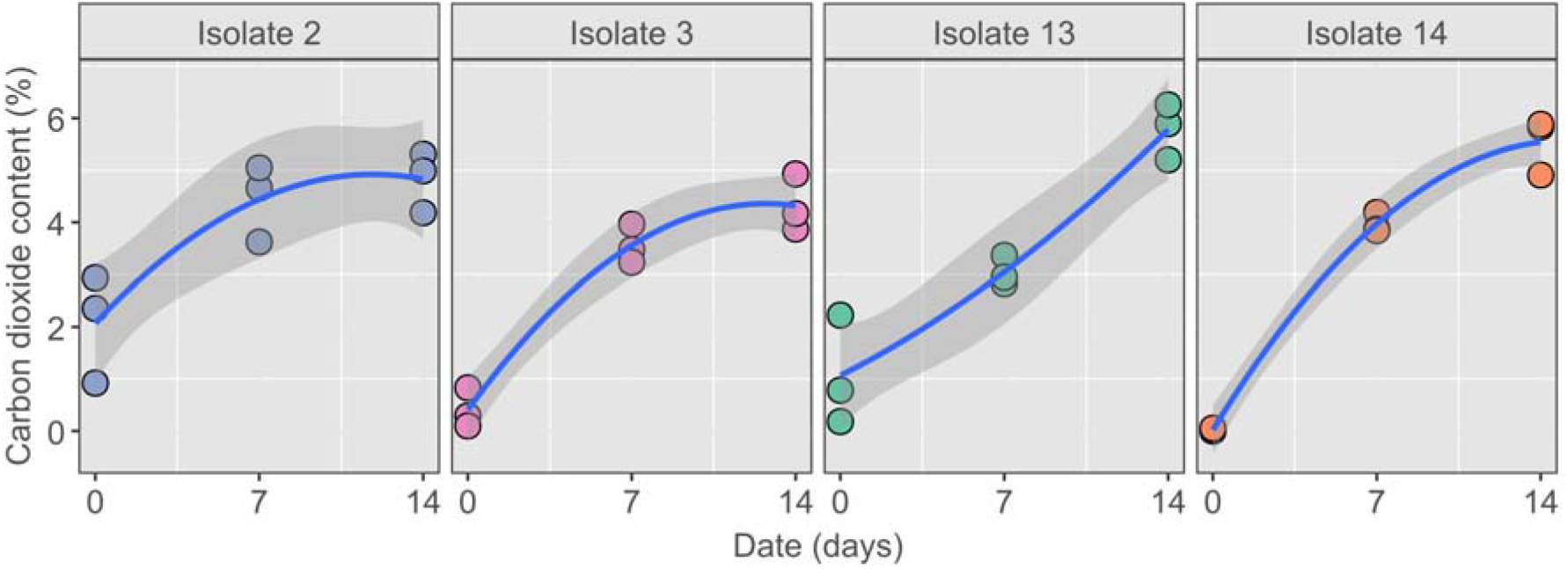
CO_2_ production (%) was measured at day 0, 7 and 14. Fully grown bacterial suspensions were used at day 0. TSB and water were used as controls (not shown). Isolate 2 and 14 produced the highest percentage of CO_2_ in the first 7 days, although Isolate 13 caught up after 14 days. The best-fit curves indicate that CO_2_ production by isolates 2, 3, and 14 follows a logarithmic growth pattern, rising rapidly on the first days. The shaded area around th fitting curves shows 95% confidence interval.

### Adaptive laboratory evolution for high pH tolerance

Given that lime has a pH of 12.4 in its freshest state, it can be a strenuous environment for bacteria to survive. Hence, an adaptive laboratory evolution experiment was performed to test whether it was possible to improve a strain’s tolerance to the high pH and if this could produce a more suitable strain to use in the lime environment. *S. clausii* was chosen as it grew faster than *S. patagoniensis* at pH 11 (Figure S 4) and it was hypothesized that a faster-growing strain is more suitable to use in a high pH environment. *S. clausii* was grown repeatedly in TSB pH 11 for 10 transfers. Initial tests were done by mixing lime with TSB but the medium would immediately precipitate, likely from the protein insolubilizing from the extreme pH range. Adjusting TSB with only NaOH was discarded due to the rapid drop in pH from a non-buffered system. Finally, TSB was adjusted to a pH of 11 using a CAPS buffer, which was the Good’s buffer with the highest pK_a_. The time for *S. clausii* to reach maximum OD was around 36 h (not shown), instead of the 48 h of Figure S 4 and Figure S 5, despite the same conditions. This was because the starting inoculum was a 1:100 dilution from a fully grown stock, rather than an inoculum with OD_600_ of 0.01 as used for the experiments shown in Figure S 4 and Figure S 5. The purpose here was to rapidly adapt the strain to the high pH instead of comparing growth kinetics. After 10 transfers the estimated number of generations was 34. A stock from the last transfer alongside the original stock was then grown in parallel in a second experiment. Gompertz curve fitting showed that the adapted strain had an increase in growth rate although it was not significant (*p* > 0.05) (Figure S 5). This newly adapted strain was deposited at the BCCM (LMG 33236, GenBank Accession No. PV163963) and is referred to as *S. clausii* transfer 10 (*t10*).

### Metabolic kinetics in a high pH environment

After the selection and adaptation of the isolated strains, a more comprehensive study was performed to study the response of the strains to a high pH. This was performed in serum bottles with TSB adjusted to pH 11 and the consumption of O_2_ and production of CO_2_, alongside the media’s pH and the cell concentration, were followed. Fully grown bacterial suspensions were used for each isolate, and along the rest of the experiments, and these were centrifuged and resuspended in fresh medium prior to the experiment.

All the strains showed a logarithmic consumption of O_2_, and *S. patagoniensis* was the fastest consumer based on the regression model, albeit not significantly (*p* > 0.05) (Figure 3). For all strains, the O_2_ levels rapidly declined within 7 days to a minimum baseline of 7.22 ± 0.2 %. To test the bacterial requirements for O_2_, at day 7 and once O_2_ consumption had seemingly stopped, two bottles from each condition were briefly opened and the internal atmosphere was equilibrated with the external atmosphere (Figure 3 gre outlined triangles). This resulted in an increase in O_2_ levels which were then rapidly reduced after three days. This shows that the system approached anoxic conditions. Nevertheless, when O_2_ became available, the bacteria were still capable of increasing their metabolism again.

**Figure 3.**
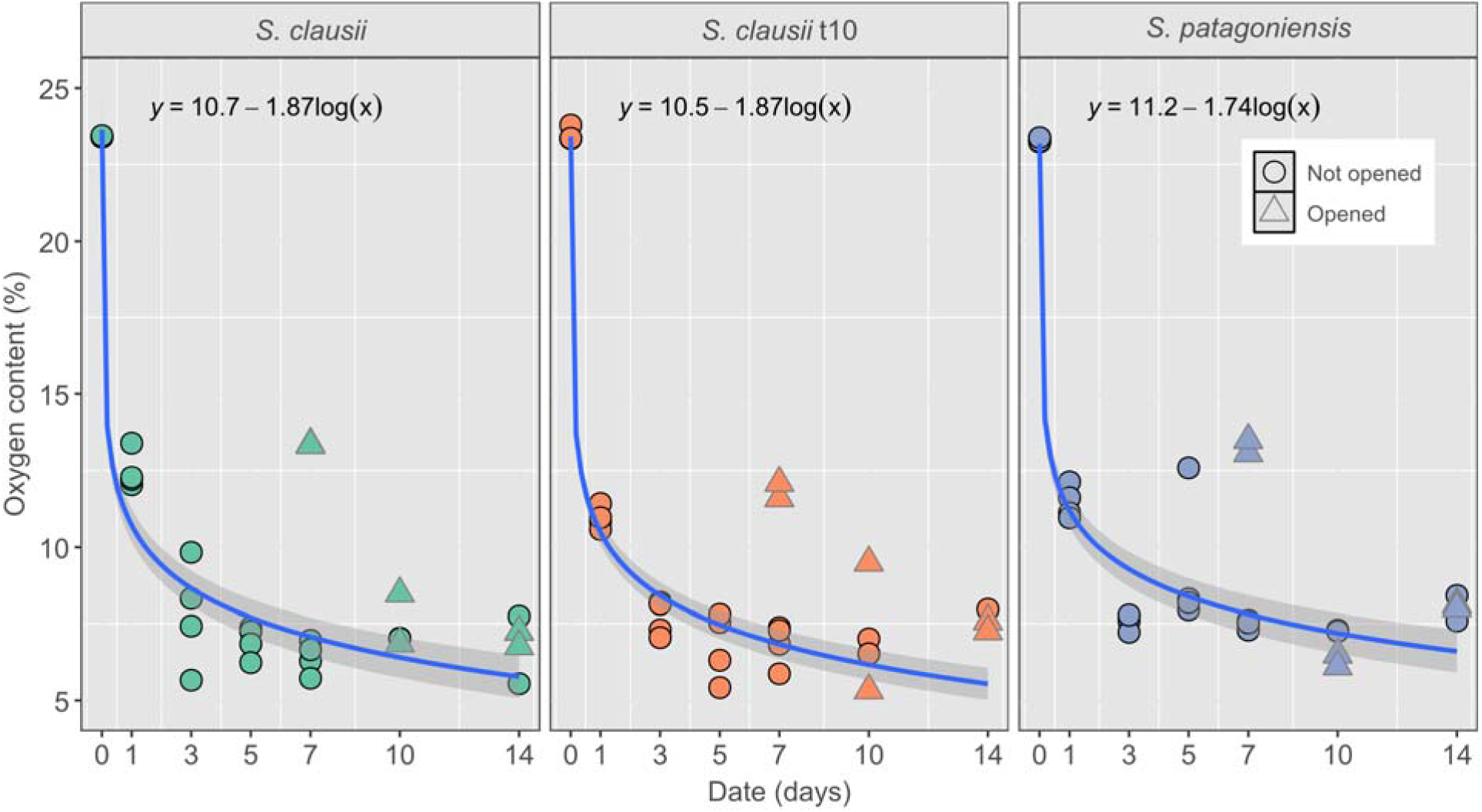
O_2_ content in the headspace of the serum bottles over time of the three tested bacteria, S. clausii, th adapted S. clausii and S. patagoniensis. Fully grown bacterial suspensions were used at day 0. Two bottles wer briefly opened at day 7 (triangles with grey outline) which allowed the O_2_ content to increase but O_2_ was readil consumed by day 14. All controls showed a much more stable O_2_ content (Figure S 3). Regressions were performed on non-opened bottles. The shaded area around the fitting curves shows 95 % confidence interval.

As expected from the consumption of O_2_, the bacteria produced CO_2_ through oxidation of organic compounds, e.g. glucose and/or amino acids in the TSB. Nevertheless, the percentage of CO_2_ was unmeasurable for most of the experimental campaign. This was attributed to the pH of the media, which acts as a CO_2_ sink. Due to the high pH, CO_2_ would first dissolve in the media and accumulate as bicarbonate and carbonate species before headspace CO_2_ concentrations, above the limit of detection, would be reached. Therefore, CO_2_ was also indirectly measured following the pH of the media, that started at 11 and by day 14 dropped to between 9 and 10 in all cases, except for the control where the pH remained at a high stable value (Figure S 6). By day 14 there was a measurable concentration of CO_2_ in the headspace of *S. patagoniensis* samples and in one *S. clausii t10* sample, but not in any *S. clausii* sample (Table 1).

**Table 1.** CO_2_ and dissolved inorganic carbon in the system at day 14 estimated from pH measurements.

| Sample<sup>1</sup> | # | Opened<br>at day 7 | m[CO<sub>2</sub>]<sub>h</sub><br>(%) | pCO<sub>2</sub><br>(atm) | [C<sub>inorganic</sub>]<br>(mmol/L) |
| --- | --- | --- | --- | --- | --- |
| *S. patagoniensis* | 4 | yes | 1.0937 | 0.0091 | 160.80 |
| *S. patagoniensis* | 3 | yes | 0.4783 | 0.0042 | 88.34 |
| *S. patagoniensis* | 2 | no | 0.2396 | 0.0013 | 93.44 |
| *S. patagoniensis* | 1 | no | 0.3884 | 0.0026 | 149.77 |
| *S. clausii t10* | 4 | yes | 0.0325 | 0.0003 | 6.87 |
| *S. clausii t10* | 3 | yes | 0.0000 | 0.0000 | 0.00 |
| *S. clausii t10* | 2 | no | 0.0000 | 0.0000 | 0.00 |
| *S. clausii t10* | 1 | no | 0.0000 | 0.0000 | 0.00 |

<sup>1</sup>No CO<sub>2</sub> was measured for *S. clausii*. Detailed calculations can be found in Table S 3.

Due to the dissolution of the CO_2_ in the media, the headspace measurements did not provide the real total amount of inorganic carbon in the system. Therefore, the CO_2_ concentration was estimated using the ideal gas law (Eq. 4 - 11) in a similar fashion as done by (33). The estimated amount of total inorganic carbon in the system by day 14 ranged from 88 to 160 mmol C/L for *S. patagoniensis* while opening or closing the bottles did not lead to a significant change (Table 1), despite that all opened bottles showed a lower pH than closed bottles for *S. patagoniensis*. The only *S. clausii t10* replicate that had a measurable amount of CO_2_ was 10 to 100 times lower (6.87 mmol C/L).

The cell concentration throughout the experiment was measured with flow cytometry at 0, 3, 7 and 14 days (Table S 2). Before the start of the experiment at day 0, all strains were grown in the same initial conditions until a fully grown bacterial suspension was achieved, and the suspension was centrifuged and resuspended in fresh media. Despite these constant initial growth conditions, at day 0, *S. patagoniensis* had a significantly higher intact cell concentration (7.4 x 10^5^ cells/ml) than the other strains (*p* < 0.05). *S. clausii t10* had more than twice the cell number than *S. clausii* (5.4 x 10^5^ vs 2.0 x 10^5^ cells/ml, respectively) and the difference was significant. This was the main observable difference between the adapted and non-adapted strains. In the first 7 days, the intact cell population had little variation but by day 14 there was a drop in cell concentration in all cases (2.3 x 10^5^, 1.8 x 10^5^ and 6.1 x 10^4^ cells/ml for *S. patagoniensis*, *S. clausii t10* and *S. clausii,* respectively). Through the whole experiment, *S. patagoniensis* and *S. clausii* t10 had significantly higher amount of intact cells than *S. clausii* (*p* < 0.05). On the other hand, the damaged population experienced a sharp drop by day 3 and then increased until day 14. In both these cases, *S. patagoniensis* had a higher damaged cell population than the other strains (*p* < 0.05). The drop from day 0 to day 3 is likely due to all the initial damaged cell populations from the resuspended bacteria continuing to break down into debris, no longer being measurable at day 3. Then, as O_2_ got depleted (Figure 3), and the fresh media got spent, the intact cells began to reduce and the damaged cell count increased (Table S 2). We can see the exact opposite effect for the opened bottles, where there was a rise in cell count at day 14 for *S. clausii t10* and *S. clausii* (open: 2.6 x 10^5^ and 1.5 x 10^5^ cells/ml vs. closed: 1.8 x 10^5^ and 6.1 x 10^4^ cells/ml, respectively) likely from the increase in metabolism due to O_2_.

Opening the bottles led to a decrease in intact cells for *S. patagoniensis* (open: 1.7 x 10^5^ vs. closed: 2.3 x 10^5^ cells/ml at day 14) (Table S 2). Interestingly, there was a larger increase of damaged cells observed for *S. patagoniensis* in the opened bottles than in the closed ones (open: 3.9 x 10^5^ vs. closed: 2.4 x 10^5^ cells/ml). This likely means that the increase in damaged cells was due to a rapid recovery of the cell population, where the intact population began to replicate, reaching a stationary phase and finally a death phase where cells began to degrade, all between day 7 and 14. This increase was not measured in the intact cell count at day 14 due to the time between measuring steps, but can be inferred from the flow cytometry density plots at day 14 by comparing intact and damaged populations (Figure S 7). The damaged cells in the plots of opened samples are denser and clustered together, suggesting a recent damage, and likely death, of the cells as the cells are still relatively intact. In contrast, the closed samples had a long cloud of red fluorescence, extending to the background, associated with different degrees of cell degradation and debris. This could further explain why the opened bottles of *S. clausii* did not have an increase in damaged cells unlike its intact cells. Given the overall lower rate of O_2_ consumption and CO_2_ production from *S. clausii* we can assume that the replication rate for *S. clausii* is slower and, while *S. patagoniensis* cells have replicated and died, *S. clausii* are only just replicating. As such, *S. clausii t10* replicates faster than *S. clausii,* but slower than *S. patagoniensis*.

The O_2_ consumption per cell, or specific rate of O_2_ consumption, (Figure 4) was calculated to determine the metabolic activity of each species (27). On average, *S. clausii* is the biggest O_2_ consumer, most starkly observed in the first days of the experiment (*p* < 0.05). By day 7, *S. clausii t10* and *S. patagoniensis* are no longer consuming O_2_, while *S. clausii* still shows activity. Overall, high specific rate of O_2_ consumption is associated with higher metabolic rates, and it is possible that the *S. clausii*, as it has less adapted to the high pH environment, is spending more energy. This could also explain the limited total cell numbers, as it is not in an optimal environment. For all strains, opening the samples at day 7 led to an increase in O_2_ consumption per cell by day 14.

**Figure 4.**
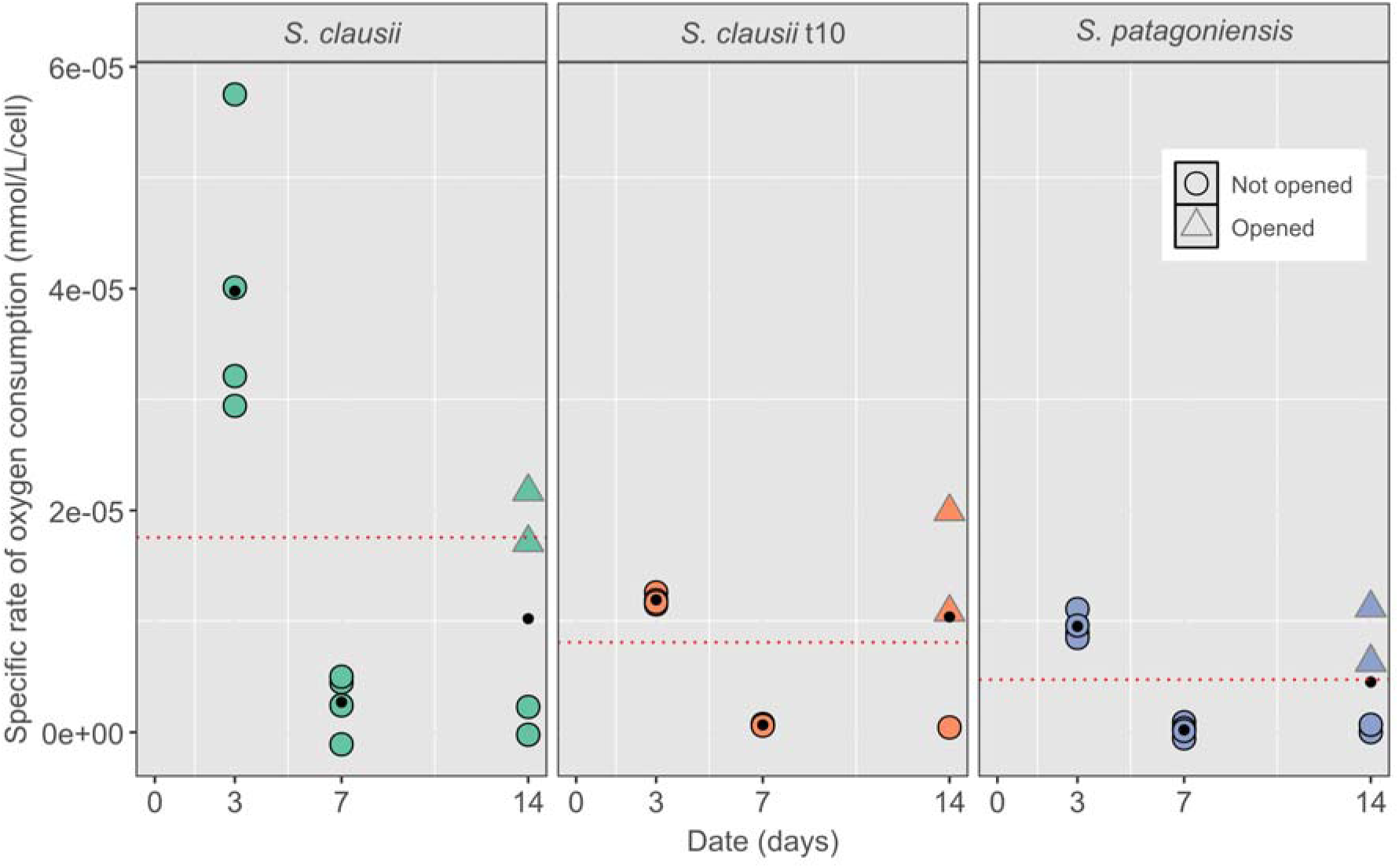
The specific rate of O_2_ consumption (mmol/L/cell) was calculated to observe the general O_2_ consumption of each species. O_2_ consumption can be directly related to the production of CO_2_. Triangles with grey outline indicate bottles opened at day 7. Black dots indicate average for the date. Red dotted line indicates average for each species and S. clausii had significantly higher consumption than the other two strains (p < 0.05).

### Carbonation of lime mortar samples

Although isolates were selected for their CO_2_ producing capacity, it was necessary to test whether carbonation of lime was possible with the bacterial suspensions. To do so, lime mortar samples were carbonated in an enclosed environment with cotton soaked in fully grown bacterial suspensions in fresh media from each isolate. The samples were incubated for 7 days at which point the experiment was terminated. All samples had a baseline of carbonation as seen on all three controls (Figure 5A), likel arising from the preparation process of the mortar samples. Nevertheless, the *S. patagoniensis* suspension seemed to have further carbonated the samples to a higher degree than the controls of air, water and TSB, although this difference was not significant (*p* > 0.05).

**Figure 5.**
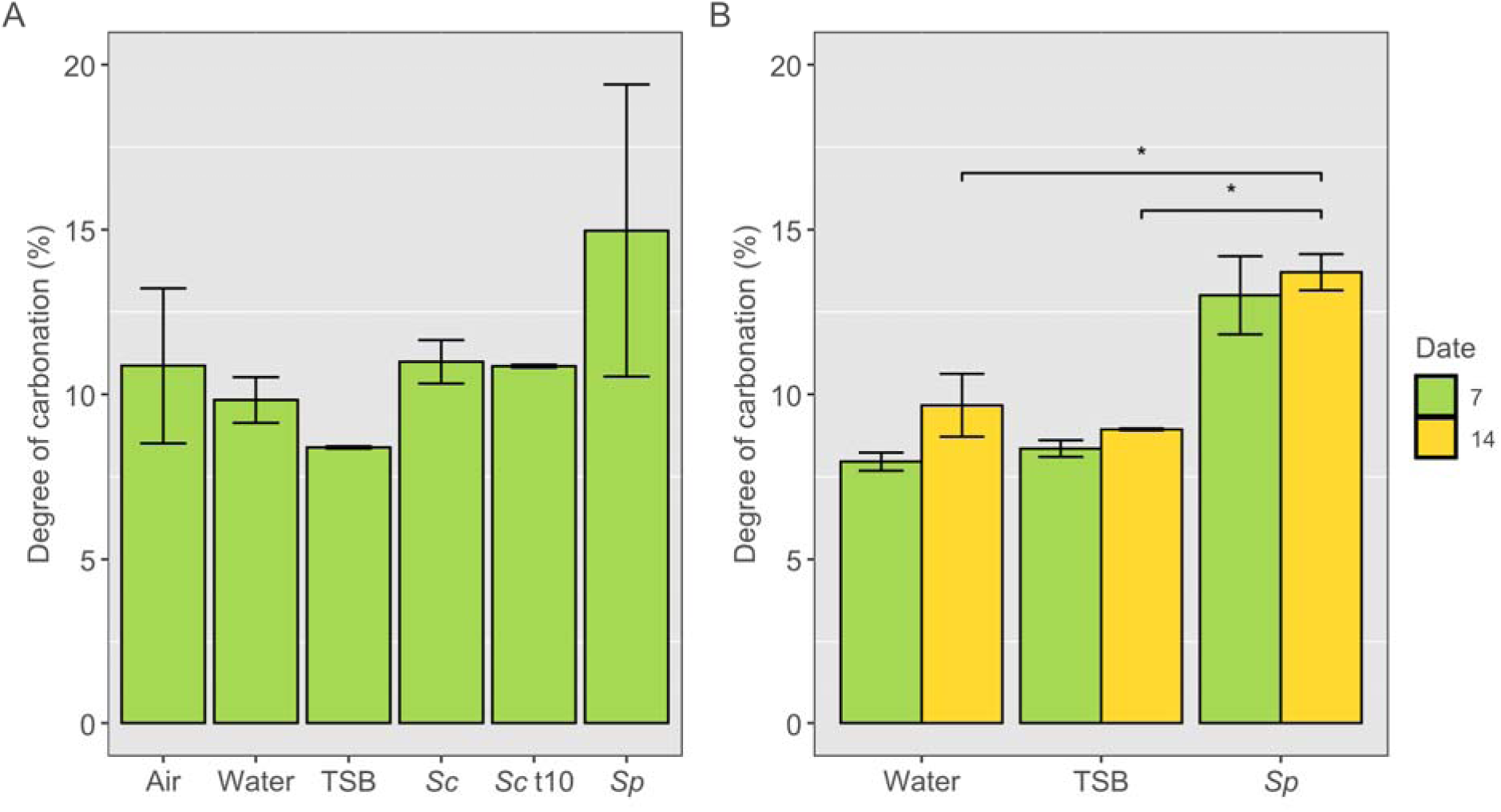
Degree of carbonation of lime mortar samples for different suspensions tested up to 7 days (n = 4) (A) an replicate tested up to 14 days (n = 6) (B). The cottons were soaked with water (Water), TSB (TSB), S. clausii (Sc), S. clausii t10 (Sct10) and S. patagoniensis (Sp). * indicates significantly different (p < 0.05). Error bars indicate standar error. Thermogravimetric curves can be found in Figure S 7. No significant different was observed between the conditions (p > 0.05).

To increase confidence in the effect of bacterial carbonation results, the experiment was repeated using *S. patagoniensis* and extended to 14 days. Once again, a baseline carbonation is observed on the Water and TSB controls, and *S. patagoniensis* also led to a significant higher carbonation than both (*p* < 0.05) (Figure 5B). By 14 days no further progression of carbonation seems to have taken place in the system, indicating a saturation of the carbonation capacity. A variability in this carbonation capacity was also observed, as measured by the larger standard error of the bacterial carbonated samples compared to the controls.

### Bacterial-lime pastes

*S. patagoniensis* and *S. clausii* t10 were chosen to perform experiments in which they were directly mixed into lime pastes. Fully grown bacterial suspensions were used for both, and these were centrifuged and resuspended in fresh TSB-C. While *S. patagoniensis* was the best-performing strain from the results of the carbonation of the lime mortar samples, *S. clausii t10* was seen to grow better at high pH and was therefore also examined. The evolution of the carbonation in the pastes was followed by TGA, FTIR and phenolphthalein spraying and the lime pastes were tested at 7, 12, 15, 21 and 35 days (Figure 8). Pastes were also prepared by mixing lime with water and TSB-C to act as controls and determine the effect of bacteria addition. Previous work has shown that killed bacteria mixed with mortar do not affect carbonation and such control was therefore not performed in this experiment (16).

TGA indicated that the water-based pastes achieved the highest carbonation percentage (Figure 6A). It followed a logarithmic relationship to time, as expected in these systems (10). The other pastes – TSB-C paste, *S. clausii* t10 paste and *S. patagoniensis* paste *–* instead did not, but carbonated linearly with respect to time, indicating a different carbonation process (Figure 6A). Overall, TSB-C pastes carbonated faster than the bacterial pastes at the beginning, matching the rate of the water pastes controls at day 7, but by 35 days the bacterial pastes had an increased rate of carbonation, matching the carbonation degree of TSB-C pastes. The TGA curves for the water pastes control show the expected behaviour, with predominant mass losses between 350 – 550 °C (corresponding to portlandite) and from 600 – 950 °C (corresponding to calcium carbonate). TSB-C and bacterial pastes showed a third mass loss between 25 – 200 °C, likely from organic matter (Figure S 9A).

**Figure 6.**
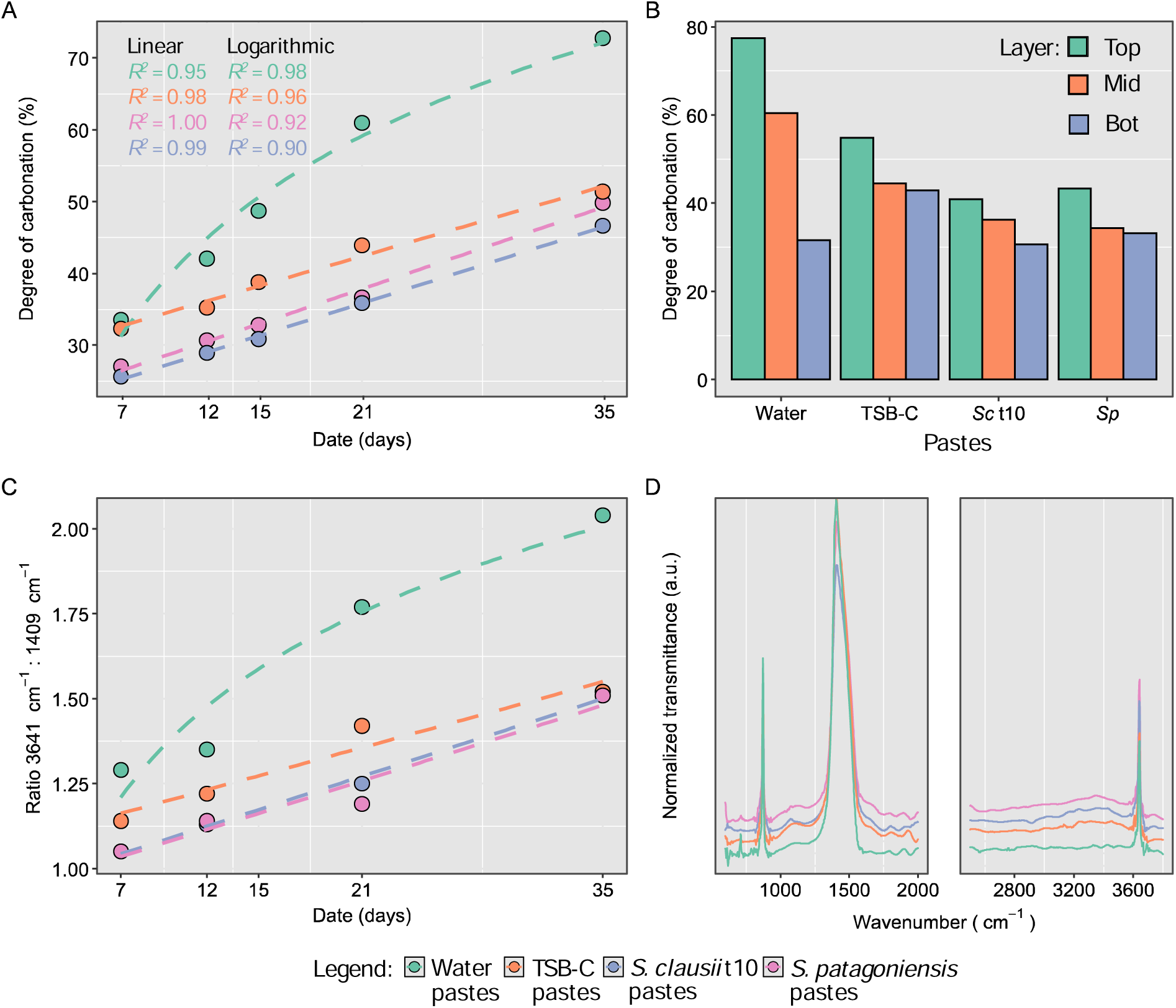
Lime paste samples with water (green), TSB-C (orange), S. clausii t10 (blue) and S. patagoniensis (purple) were tested at 7, 15, 14, 21 and 35 days using TGA, FTIR and phenolphthalein analysis. A) Degree of carbonation of lime pates determined by TGA. R^2^ values for linear and logarithmic regressions of the progression of carbonation of all pastes. TSB-C and bacterial pastes have a better fit with linear models while water’s best fit is logarithmic. TGA curves can be found in Figure S 8A. B) Degree of carbonation for the separate layers of the sample top (blue), mid (orange), green (bottom). TGA curves can be found in Figure S 8B. C) Ratio between the absorbance of OH band (3641 cm^-1^) of Ca(OH)_2_ and the v_3_ mode of CaCO_3_ at (1409 cm^-1^). D) FTIR spectra of the lime pastes at day 15 between 600 – 2000 cm^-1^ and 2700 - 3700 cm^-1^, where organic signatures were detected. Given that one sample per paste and date was tested, no statistical analyses were performed.

At day 21, the pastes were separated into 3 layers (top, mid and bottom) and grounded to evaluate the progression of carbonation throughout the sample’s profile (Figure 6B). In all cases, the top layer was the most carbonated. In the pastes with water, the mid layer was also more carbonated than the bottom layer. For TSB-C and bacterial pastes, these two lower layers were similarly carbonated. Remarkably, for the TSB-C paste, the bottom layer had more carbonation than that of the water paste, showing that carbonation progressed more evenly on these samples than the control. For the bacterial pastes, the bottom layer was equally carbonated as the pastes made with water only, meaning no effect was measurable. All TSB-C and bacterial layers also showed a weight loss during TGA between 25 – 250 °C, coming from the presence of organic material (Figure S 9B).

FTIR confirms the presence of organic phases, present as a shoulder between 1500 and 1700 cm^-1^ and as a broad band between 3000 – 3600 cm^-1^ (Figure 6D). These were ubiquitous in TSB-C and bacteria containing pastes. Moreover, in all cases characteristic *v*-modes of calcite were observed (712, 874 and 1409 cm^-1^) and a strong OH stretching (3641 cm^-1^), corresponding to portlandite. Interestingly, the development over time of the ratio between the portlandite peak vs the main calcite band (1409 cm^-1^) was remarkably similar to that of the percentage of carbonation determined using TGA (Figure 6C). The v_3_-mode has been used to determine carbonation extent in cement-based materials (34).

A standard progression of carbonation was observed following phenolphthalein application for pastes with water, where a whiter (more carbonated) region progressed downwards to the bottom, less carbonated (pink) region of the sample, along the direction of carbonation (Figure S 10). Instead, for TSB-C and bacterial pastes a two-step carbonation progression was observable. Here, a slow drop in pH of the lime pastes occurred, whereby at 7 days the carbonation front was not visible. At 12 days whiter regions were observable in the TSB-C and bacterial pastes, but only at day 15 was there a visible white band. This carbonated region was centered in the sample, in between two pink bands of uncarbonated lime, a thin one on the surface and a thicker at the bottom, showing that the carbonation front progressed in a non-uniform, discontinuous way (Figure S 10). Conversely, when the samples were tested at day 21, we observed that the carbonation front was measurable again and that the surface band of dark pink disappeared. At day 35 this phenomenon repeated, a lighter center indicating carbonation was surrounded by pink bands, although slight variations per samples were visible. This phenomenon was observable when the experiments were repeated, although different rates of progression were detected.

## Discussion

The focus of this work was to investigate how certain bacteria could produce CO_2_ to speed up the carbonation process of lime (Eq. 1) and to produce a more thorough carbonation, which would help lime mortars set faster at an early age. Due to the high pH of lime (12.4 in fresh state), only alkaliphilic bacteria were deemed suitable to use here. An isolation campaign was conducted from an old lime mortar in a masonry wall, and isolates were selected based on their alkalophilicity and CO_2_ production capacity, while further characterization of their O_2_ consumption rates and cell numbers was performed. Finally, the capacity for bacterial CO_2_ to carbonate lime mortars was tested and the impact of mixing bacterial suspensions with lime on the carbonation process was examined.

Thirty-three isolates were isolated from a lime mortar wall and were characterized through MALDI-TOF and sequencing to obtain sixteen distinct strains. All genus results were expected given the type of sample and were mostly alkaliphilic. The most common ones – *Shouchella*, *Bacillus* and *Staphylococcus –* are all part of the Firmicutes phyla, which is dominant in limestone environments (35, 36). *Shouchella* are one of the main Bacillus clades and are particularly known for being alkaliphiles (37). Four isolates showed a capacity to grow fast and effectively at pH 11, which is one of the highest pHs achieved with Good’s buffer solutions. From these 4 isolates, 3 were fast CO_2_ producers and the best two were selected for further study, namely an *S. clausii* and *S. patagoniensis* strains. *S. clausii* was further adapted to a high pH environment in an adaptive laboratory evolution experiment. *S. clausii* was chosen over *S. patagoniensis* due to its better growth kinetics, under the hypothesis that a faster growth rate means better adaptation to high pH and, thus, better suitability to use in lime. The isolate was grown over 10 transfers in TSB pH 11, and a total of around 34 generations. The adapted strain, *S. clausii* t10, was observed to have a faster growth rate than *S. clausii*, albeit not significant, but had a significantly higher stationary phase cell density at pH 11, observed through cell counts. We consider the higher cell counts for *S. clausii* t10 an indicative of low stress. This is because more energy can be invested into replication rather than the high energy needs for cells to maintain homeostasis or other non-growth-associated metabolic activities due to suboptimal environment (32). In turn, this leads to an increase in biomass at stationary phase (32, 38, 39), coinciding with what is observed in the results presented. Despite the other limited results, the higher cell count was an indicator that the adaptive laboratory evolution of *S. clausii* was partly successful. The number of generations, 34, is on the lower-end of what is common in adaptive laboratory evolution (40), and extending the length of the experiment would have possibly provided a better result.

All three strains, *S. clausii, S. clausii* t10 and *S. patagoniensis,* were then studied in depth in a parallel experiment, following their growth dynamics and capacity for carbonation of lime mortars. In this experiment, a fully grown suspension was used as replicating bacteria have a lower catabolic activity and will produce less CO_2_ than those in stationary phase (32). In stationary phase the cells are not replicating, biosynthetic pathways are less active, and C is not incorporated into macromolecules, and catabolic activity is higher. Given that the objective is to achieve better carbonation, where higher CO_2_ concentrations are desired, then catabolic activity was preferable. The use of a fully grown suspension also ensured that there was no contamination or competition by possible indigenous species, due to the high cell density outcompeting any possible colonization of the nutrients. The experimental series following metabolic kinetics in serum bottles showed that *S. patagoniensis* had the highest intact cell count of all strains when grown in TSB pH 11, and had a high replication rate between day 7 and 14. Moreover, *S. patagoniensis* was the fastest consumer of the O_2_ in the headspace of the bottles. Even when the samples were opened on day 7 to introduce more OL, *S. patagoniensis* was the quickest to increase its metabolism and deplete the available OL, reaching a second stationary phase ahead of the other two strains. Additionally, *S. patagoniensis* exhibited the lowest specific rate of OL consumption. Since the specific rate reflects metabolic activity, a lower demand for OL indicates a lower organic acid consumption, i.e. lower energy demand (41, 42). This lower energy demand could be attributed to a better adaptation to the high pH environment. Overall, *S. patagoniensis* seemed the best adapted to the high pH outperforming *S. clausii* and the adapted *S. clausii* t10. Lastly, *S. patagoniensis* was the only strain that produced a measurable amount of CO_2_ in the headspace of the serum bottles, which confirms that it was the best CO_2_ producer of the studied strains. Unfortunately, CO_2_ in the headspace could only be measured at day 14, and the evolution of CO_2_ production could not be followed. This was due to the high pH environment in which tests were conducted which acted as a carbon sink and it was only after pH reached 9 – 9.5 that CO_2_ began to accumulate in the headspace. Nevertheless, it is possible to correlate the aforementioned cell density and growth phase, as well as inversely correlate the consumption of O_2_ of *S. patagoniensis* with the production of CO_2_ (32).

The specific rate of O_2_ consumption, also allows to compare species in different starting conditions (43). Here, the cell count differed per species, but the starting concentration in each experiment was not altered as species-specific differences in similar starting conditions were of interest in determining the most suitable strain. Through the use of O_2_ consumption per cell it is possible to see that the higher cell count of *S. patagoniensis* accounted for the larger CO_2_ production of all three strains. This contradicts our initial hypothesis that a more adapted strain will more readily enter catabolic metabolism, but it is possible that the high stress on *S. clausii* is leading to an even higher increase in activity of these catabolic pathways as more energy is needed to maintain homeostasis. Furthermore, this also means that the least adapted strain will have a lower production but a higher CO_2_ yield (i.e. the production per unit). As such, depending on the optimal concentration of cells needed to carbonate lime, a different strategy to that hypothesized here might become more suitable: if a lower cell count is possible, a less adapted strain with a higher CO_2_ yield would be a more optimal carbonation solution.

The measurements of CO_2_ at day 14 also allowed to estimate the total amount of CO_2_ produced by the bacteria that was present as CO_2_ in the headspace, CO_2_ dissolved in the media, and CO_2_ converted to H_2_CO_3_, HCO_3_^-^ and CO_3_^2-^. The amount of CO_2_ produced by *S. patagoniensis* by day 14 was 88 to 160 mmol/L. This accounts to a CO_2_ production rate of 6 to 12 mmol/L/day, or 6.94 x 10^-5^ to 1.39 x 10^-4^ mmol/L/s (assuming a constant production rate through the whole experiment which is not the case). Following Van Balen and Van Gemert where it is assumed that all CO_2_ will react with portlandite in a mix system (with excess Ca(OH)_2_), the production rate of CO_2_ by the bacteria becomes comparable to the values reported by these authors for CO_2_ uptake during carbonation (6). In this case, the upper limit of *S. patagoniensis* production capacity is relatively close to the lowest margin of the maximum CO_2_ uptake by lime in an ideal scenario (3.99 x 10^-4^ 1/s). Given that no optimization for maximizing CO_2_ production was performed on *S. patagoniensis*, and that these values are conservative, we can conclude that the use of bacterial CO_2_ might provide a viable and useful method for early age carbonation of lime. Lastly, when looking at the carbonation of the lime mortar samples in a closed atmosphere, *S. patagoniensis* carbonated the samples faster and to a higher degree than the other two strains. Still, in the case of lime mortar samples, a high variability was observed between runs and the carbonation did not progress further after 14 days compared to 7 days. This was not the case in the serum bottle experiment, where CO_2_ was highest at 14 days. This is expected to be due to the different setups with which carbonation and CO_2_ measurements were tested.

To show the possibility of bacteria to be used as an earlier hardening agent of lime, we performed direct mixing of the bacteria suspensions with lime in pastes. The use of pastes allows to confirm whether the changes in the carbonation progress are more directly attributed to the bacterial additives and no other interactions that can be caused by aggregates (44). Interestingly, a sort of inhibitory effect was observed for the TSB-C and bacterial pastes when compared to the water controls, but a very different carbonation dynamic was observed. A two-step carbonation process was observed, whereby a lower pH (linked to the carbonation front) could be measured only after 12 to 15 days, depending on the mixture. Still, this region of carbonation was sandwiched between two uncarbonated bands closer to the surface and bottom of the sample, respectively. To explain the advance of the carbonation front past an uncarbonated region, we hypothesize that it was due to the development of a Liesegang pattern (44, 45). This type of pattern appears in porous systems subjected to diffusion-reaction processes, as it is the case of a lime paste undergoing carbonation. The diffusion (porous) medium is the lime paste itself and the diffusing reactant is CO_2_ towards the uncarbonated portlandite particles. Following carbonation, an amorphous calcium carbonate (ACC) metastable nanophase forms which can also diffuse in a partially saturated medium and eventually transform into a stable crystalline CaCO_3_ (calcite) at a particular location where a Liesegang band appears (i.e., lighter band surrounded by pink bands after phenolphthalein application). After this event, the carbonation front progresses to the sample core where another precipitation event can take place forming another CaCO_3_ band (45). However, the dissolution of uncarbonated portlandite and redistribution of this solution along the pore space of the lime plaster can lead to a pH increase even in areas where carbonation has already taken place, resulting in the periodic bands associated with the Liesegang phenomenon, usually formed in rapidly carbonating samples (3). This would explain the more homogeneous carbonation observed in the different layers of the pastes mixed TSB-C tested at 21 days, whereas the pastes mixed with water had a stratified carbonation. Interestingly, these Liesegang patterns have been associated with superior lime mortars (45). The change in carbonation progression in the TSB-C and bacterial pastes can certainly be attributed to organic phases found in these mixtures. Organic presence confirmed by weight loss in the 25 – 250 °C T range using TGA and in the FTIR analysis, at the bands between 1500 and 1700 cm^-1^ and 3000 – 3600 cm^-1^.

Despite the promising effect of the Liesegang pattern formation seemingly induced by the organics, the overall carbonation was inhibited in these mortars. The use of TSB as a bacterial medium could have partly influenced this as it contains a concentration of 0.5 % of NaCl. At higher concentrations (3.5 – 7 %) NaCl rapidly increases carbonation at an early age. This creates a dense carbonation layer at the surface that inhibits the carbonation process by 28 days in aerial mortars (46). On the other hand, TSB also contains glucose, which also has an inhibitory effect on carbonation, but this leads to the opposite effect in the long term compared with NaCl, where CO_2_ can diffuse through the whole sample carbonating better inside and improve the mechanical properties of the materials (46). As such, TSB contains two compounds, although at low concentrations, that inhibit carbonation in the short term (28 days), which would explain the lower performance of the TSB-C and bacterial pastes compared to water. Finally, the bacterial biomass seems to further inhibit carbonation early on, but a ramp up is experienced by day 35, achieving similar degrees of carbonation as the TSB-C control. It is possible that bacterial CO_2_ allowed for this improvement. Nevertheless, the length of the experiment did not allow to observe any discernible bacterial effect and the mechanistic explanation for this rate improvement at the end of the experiment is not possible to outline here.

This work was not an exhaustive analysis of the bacterial strains isolated nor the results optimal. Nevertheless, it serves as a proof-of-concept for the use of bacterial CO_2_ as a strategy to improve carbonation in lime mixtures. Only two previous works have explored this possibility before, showing the topic is still in its infancy (15, 16). Importantly, our work expands on this previous research by looking in depth at a more robust selection process for isolates and correlates the production of CO_2_ with the carbonation of lime. Still, further work is required to prepare lime mortar formulations optimized for bacterial carbonation strategies. For example, the mechanisms of action of how carbonation improves need to be more rigorously studied. Here, dissolved inorganic carbon (DIC) was not measured and such data would prove valuable to confirm that the drop in pH is only associated with respiration of carbon sources and not the influence of other organic acids. Moreover, the media composition should be carefully reassessed. In this work, TSB was used, which affected carbonation when mixed directly with lime, even though the buffering component was removed (TSB-C). Moreover, TSB-C had NaCl, which in future applications could lead to salt damage (47). TSB can also be too expensive at economies of scale, and the choice of media is central in determining the scalability of a product (48), while certain media components, particularly carbon sources like glucose, can have varying carbon footprints which need to be carefully assessed (49). Finally, further research should also focus on analyzing the mechanical and physico-chemical effect that the bacterial suspensions have on the ultimate performance of the lime mortars. Tests are required to determine whether, even with changes in carbonation, other durability and microstructural properties are influenced; and compare this novel strategy to the most promising approaches to speed up carbonation of lime-based binders, such as the use of carbonic anhydrase and MOFs (50). Note that the possible formation of carbonate-organic hybrids with superior physical-mechanical properties as in the case of carbonate biominerals in general (e.g. sea shells (51)) and bacterial biominerals in particular (24), would lead to better performance not only in terms of carbonation speed at early ages, but also on improved long-term durability of these bio-based lime mortars and plaster.

## Conclusion

This study was a proof-of-concept of the potential of bacterial COL production to accelerate the carbonation of lime mortars, offering a promising bio-based strategy for enhancing early-age strength. It expands on previous studies by providing a more systematic selection of bacterial strains and directly linking COL production to carbonation efficiency. Among the isolated strains, *S. patagoniensis* exhibited the best adaptation to high pH conditions and the highest COL production, directly contributing to improved carbonation when compared to the other studied strain. Despite some promising results, challenges remain and the extent of carbonation was not optimal. Furthermore, variations in carbonation progression suggest that additional mechanistic studies are needed to fully understand the interplay between bacterial additives and lime materials.

## Author statement

**Franco Grosso Giordano:** Conceptualization, Methodology, Formal Analysis, Investigation, Writing – Original Draft. **Quinten Mariën:** Methodology, Formal Analysis. **Nele De Belie:** Funding acquisition, Writing – Review & Edition, Supervision. **Carlos Rodriguez-Navarro:** Conceptualization, Writing – Review & Edition, Supervision. **Nico Boon:** Conceptualization, Formal Analysis, Writing – Review & Edition, Funding acquisition.

## Supporting information

All supplementary data

## Acknowledgements

We thank the personnel of the Center For Microbial Ecology and Technology for the help with this manuscript. In particular to Tim Lacoere for his help with DNA extraction and sequencing of the isolates. Additional thanks go to Lhoist, in particular Ulrike Peter and Damien Grognard, for their support, help with the instrumentations and data analysis and use of their facilities. Funding was provided by the European Union’s Horizon 2020 research and innovation programme under Marie Sklodowska-Curie project SUBLime [Grant Agreement n◦955986], with co-financing by the Bijzonder Onderzoeksfond (BOF) [ITN016-22 BOF] and [BOF24/CDV/148].

