## Supplementary material for "Isolation and Characterization of Strains used in Bacterial-Based Strategies for Accelerated Carbonation of Lime Mortars": All supplementary data

**Email addresses**

### Appendix

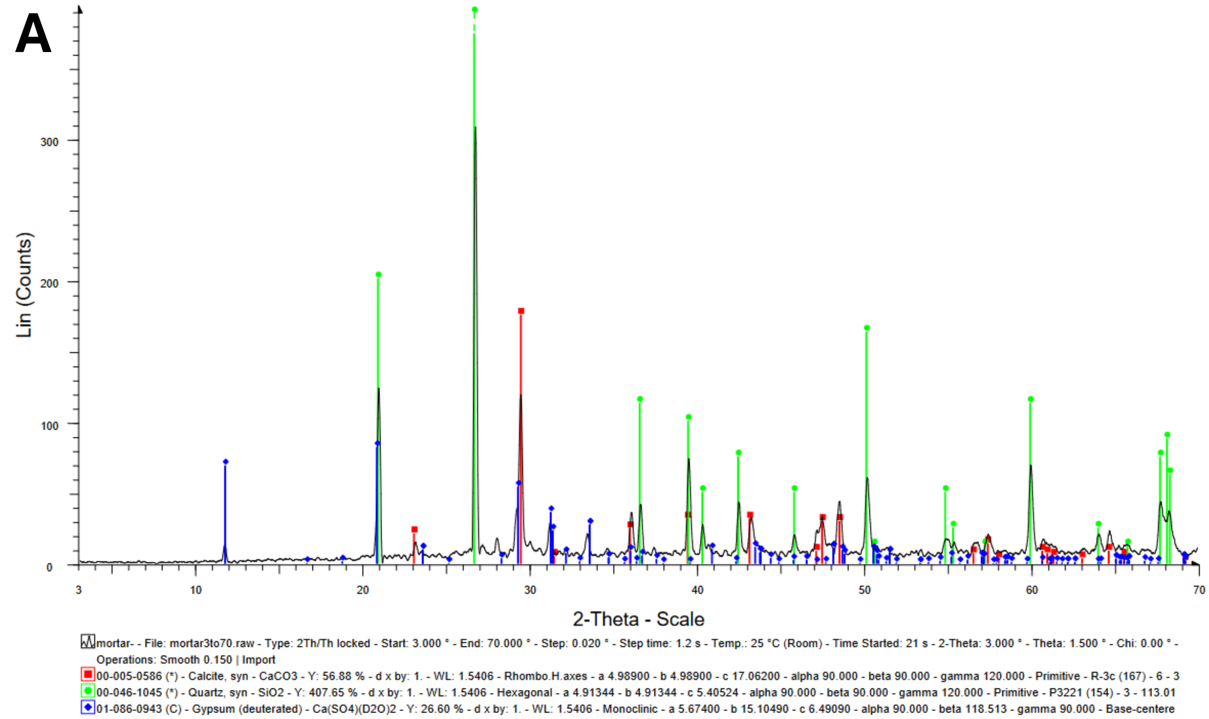
A
Lin (Counts)
2-Theta - Scale
mortar- - File: mortar3to70.raw - Type: 2Th/Th locked - Start: 3.000 ° - End: 70.000 ° - Step: 0.020 ° - Step time: 1.2 s - Temp.: 25 °C (Room) - Time Started: 21 s - 2-Theta: 3.000 ° - Theta: 1.500 ° - Chi: 0.00 ° - Operations: Smooth 0.150 | Import
00-005-0586 (*) - Calcite, syn - CaCO3 - Y: 56.88 % - d x by: 1. - WL: 1.5406 - Rhombo.H.axes - a 4.98900 - b 4.98900 - c 17.06200 - alpha 90.000 - beta 90.000 - gamma 120.000 - Primitive - R-3c (167) - 6 - 3
00-046-1045 (*) - Quartz, syn - SiO2 - Y: 407.65 % - d x by: 1. - WL: 1.5406 - Hexagonal - a 4.91344 - b 4.91344 - c 5.40524 - alpha 90.000 - beta 90.000 - gamma 120.000 - Primitive - P3221 (154) - 3 - 113.01
01-086-0943 (C) - Gypsum (deuterated) - Ca(SO4)(D2O)2 - Y: 26.60 % - d x by: 1. - WL: 1.5406 - Monoclinic - a 5.67400 - b 15.10490 - c 6.49090 - alpha 90.000 - beta 118.513 - gamma 90.000 - Base-centere

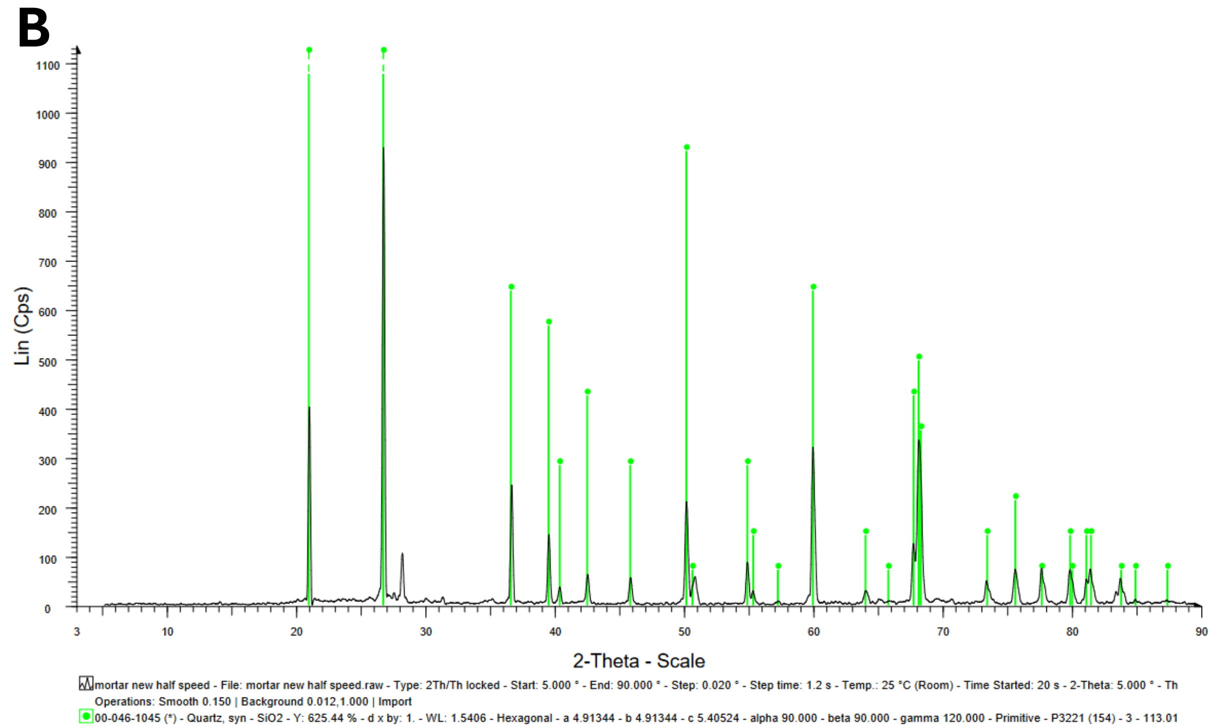
B
Lin (Cps)
2-Theta - Scale
mortar new half speed - File: mortar new half speed.raw - Type: 2Th/Th locked - Start: 5.000 ° - End: 90.000 ° - Step: 0.020 ° - Step time: 1.2 s - Temp.: 25 °C (Room) - Time Started: 20 s - 2-Theta: 5.000 ° - Th
Operations: Smooth 0.150 | Background 0.012,1.000 | Import
00-046-1045 (*) - Quartz, syn - SiO2 - Y: 625.44 % - d x by: 1. - WL: 1.5406 - Hexagonal - a 4.91344 - b 4.91344 - c 5.40524 - alpha 90.000 - beta 90.000 - gamma 120.000 - Primitive - P3221 (154) - 3 - 113.01

*Figure S 1 – XRD patterns of a fragment of lime-mortar wall before (A) and after (B) acid attack by 37 % HCl solution. All Bragg peaks in A correspond to calcite and quartz with some indications of gypsum, while only quartz and small amounts of feldspar (Bragg peak at ~28 °2θ) are observed after the acid attack (B).*

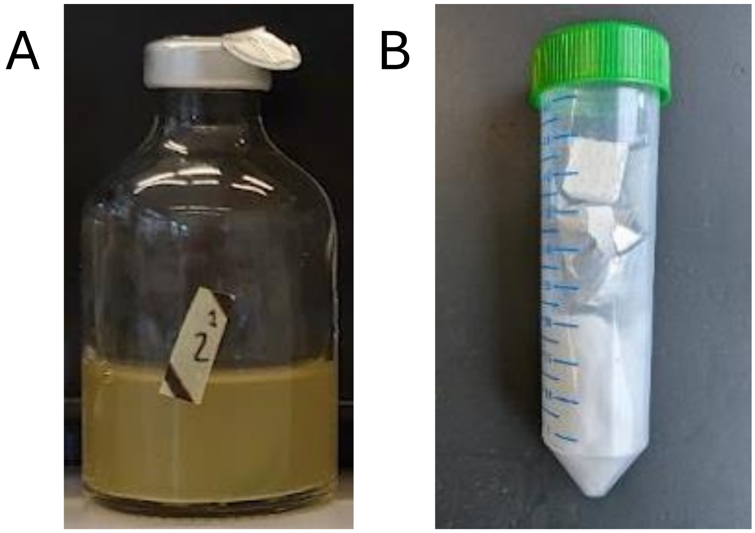


*Figure S 2 – A) 30 mL serum bottle capped with butyl rubber stoppers and sealed, used for following metabolic kinetics of the different isolates including measuring gas production, cell counts and pH over time. B) Arrangement of the soaked cotton, aluminum paper and lime mortar samples inside a Falcon tube. This system acted as a closed environment to test carbonation under an atmosphere with increased* $CO_2$ *due to bacterial respiration.*

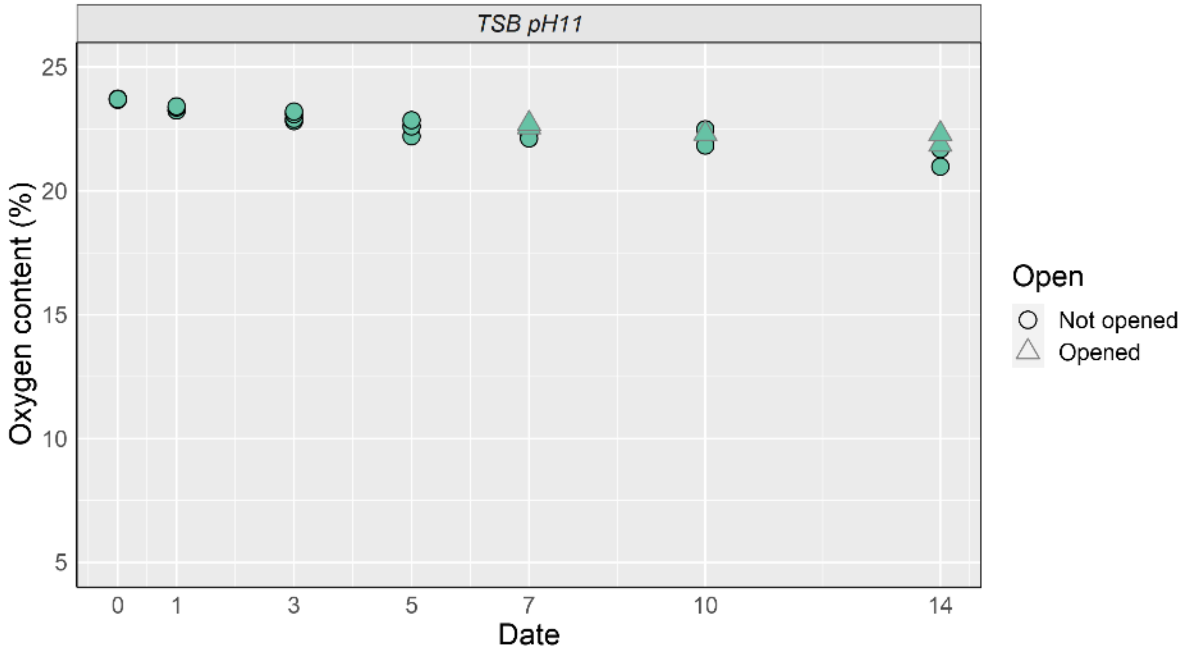


*Figure S 3 – $O_2$* content in the controls*, bottle with sterile TSB*. The $O_2$ level slowly decreases overtime *likely from sampling*. The bottles that were opened *(triangles with grey outline)* also did not see a change in $O_2$ concentration, given that the internal atmosphere remained at the same pressure so there was no active exchange of gas. *The addition of $NaN_3$ to one of the bottles shows no change in $O_2$ concentration, nor $CO_2$ (not shown).*

*Table S 1 – All isolates from the isolation campaign according to the media on which they were isolated*[1]

| Isolation media | Isolate name | Reference group | Reference strain | MALDI-TOF MS hit | Amplicon sequencing hit |
|---|---|---|---|---|---|
| R2A | R2A-3B* | A | 0 | *Bacillus sp.* | *Bacillus sp., cereus, thuringiensis, paranthracis, pacificus, toyonensis* |
| R2A | R2A-4F | A | | *NA* | |
| **R2A** | **R2A-5F** | **B** | **1** | ***Bacillus thuringiensis*** | ***NA*** |

| | | | | | |
|---|---|---|---|---|---|
| **Horikoshi** | **Hori-4D** | **C** | **2** | ***Bacillus clausii*** | ***Alkalohalobacillus clausii*** |
| **Horikoshi** | **Hori-3B** | **D** | **3** | ***Bacillus clausii*** | ***Bacillus velezensis, amyloliquefaciens, subtilis*** |
| **R2A** | **R2A-4J** | **E** | **4** | ***NA*** | ***Bacillus sp., mobilis, toyonensis, wiedmannii*** |
| R2A pH 9 | R2A9-H.B* | F | 5 | *NA* | *Bacillus mobilis, toyonensis or wiedmannii, cereus* |
| R2A | R2A-3A.A | G | | *Bacillus sp.* | |
| R2A | R2A-3A.B | G | | *Bacillus sp.* | |
| R2A | R2A-3C* | G | 7 | *Bacillus sp.* | *Bacillus thuringensis, cereus mobilis or wiedmanni* |
| R2A pH 9 | R2A9-4H.A | G | | *NA* | |
| **R2A pH 9** | **R2A9-4K** | **G** | **6** | ***Bacillus sp.*** | ***NA*** |
| **R2A pH 9** | **R2A9-3A** | **H** | **8** | ***Bacillus thuringiensis*** | ***Staphylococcus warneri*** |
| R2A | R2A-4H | I | | *Staphylococcus warneri* | |
| **R2A pH 9** | **R2A9-3C.C** | **I** | **10** | ***Staphylococcus warneri*** | ***Sphingomonas sp.*** |
| **R2A pH 9** | **R2A9-4E** | **I** | **9** | ***Staphylococcus warneri*** | ***Staphylococcus warneri*** |

| | | | | | |
|---|---|---|---|---|---|
| R2A pH 9 | R2A9-4F | I | | *Staphylococcus warneri* | |
| R2A | R2A-4E | J | | *NA* | |
| R2A pH 9 | R2A9-3C.A* | J | 11 | *NA* | *Bacillus mobilis, toyonensis, wiedmannii, cereus* |
| **R2A pH 9** | **R2A9-3C.B** | **K** | **12** | ***NA*** | ***Alkalobacillus sp, patagoniensis*** |
| **Horikoshi** | **Hori-3C** | **L** | **14** | ***NA*** | ***Alkalobacillus sp, patagoniensis*** |
| Horikoshi | Hori-4E | L | | *NA* | |
| **Horikoshi** | **Hori-5F** | **L** | **13** | ***NA*** | ***Alkalobacillus clausii*** |
| Horikoshi | Hori-5G | L | | *NA* | |
| **R2A pH 9** | **R2A9-3B** | **M** | **15** | ***Bacillus sp*** | ***Alkalobacillus sp, patagoniensis*** |
| R2A pH 9 | R2A9-5J | M | | *NA* | |
| Horikoshi | Hori-3A | N | | *NA* | |
| R2A | R2A-3D | NA | | | |
| R2A | R2A-4G | NA | | | |
| R2A | R2A-4I | NA | | | |
| R2A pH 9 | R2A9-3D | NA | | | |
| R2A pH 9 | R2A9-4G | NA | | | |
| R2A pH 9 | R2A9-5I | NA | | | |

[1]*Shown are the isolate name, the reference group according to MALDI-TOF MS analysis, the reference strain of each group according to MALDI-TOF MS analysis, the classification according to MALDI-TOF MS analysis and classification according*

*to 16S amplicon sequencing. Bold rows are the 12 candidates selected after isolation for further analysis. * indicates strains that are biosafety level 2.*

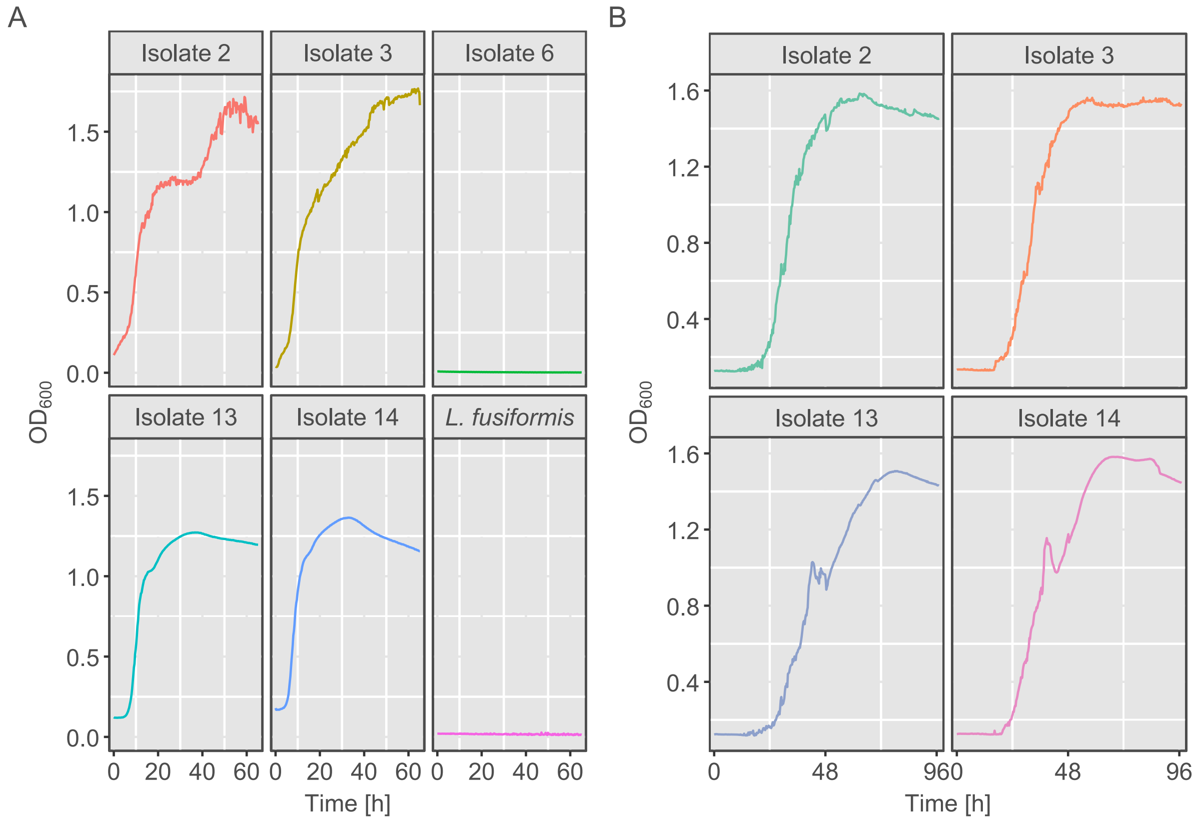


*Figure S 4 – $OD_{600}$ growth curves in TSB pH 10 (A) and TSB pH 11(B) for the best growing isolates at TSB pH 10. Isolates were inoculated from a double transferred fully grown suspension in TSB. All curves were inoculated to $OD_{600}$ = 0.01 and are average of three replicates.*

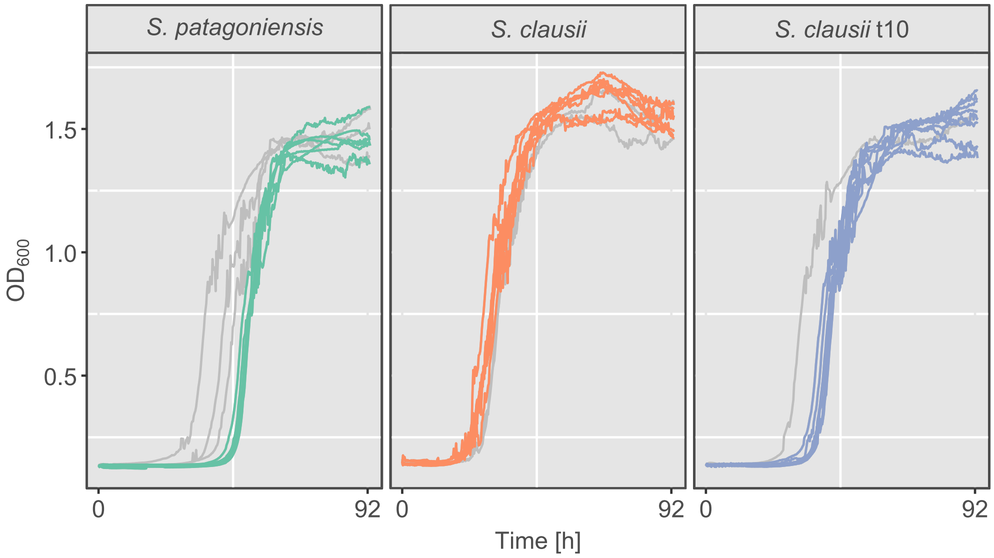


| | ***S. patagoniensis*** | ***S. clausii*** | ***S. clausii t10*** |
|---|---|---|---|
| **Max log fitted OD** | 0.16 | 0.20 | 0.17 |
| **Maximum log fitted growth rate** | 0.23 | 0.17 | 0.20 |
| **Lag phase (h)** | 43.69 | 20.19 | 34.21 |
| **MSE** | 0.01 | 0.02 | 0.02 |

*Figure S 5 – $OD_{600}$ growth curves in TSB pH 11(B) for the original strains and the ALE strain* (S. clausii *t10*). *All curves were inoculated to $OD_{600}$ = 0.01. Isolates were inoculated from a double transferred fully grown suspension in TSB pH 11. Grey outlined curves are outliers based on Iglewicz and Hoaglin's robust test for multiple outliers. The average maximum log fitted OD, growth rate, lag phase based and mean square error (MSE) on Gompertz fitting for all curves for all three tested strains. No significant difference was observed between strains* ($p > 0.05$).

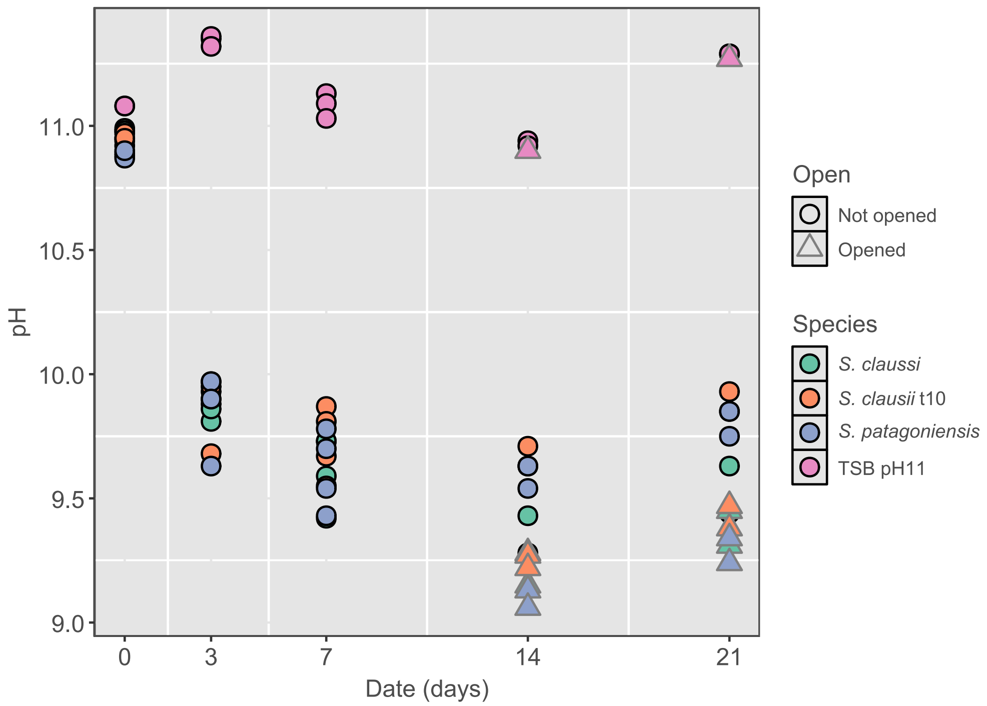


*Figure S 6 – pH measured over time for all conditions at day 0, 3, 7, 14 and 21. A steady drop in the pH is observed as $CO_2$ dissolves in the media until a steady state is achieved. No significant difference was observed between strains ($p > 0.05$). For the control, TSB pH11 (pink) no significant difference was observed ($p > 0.05$).*

*Table S 2 – Cell counts in the bacterial suspension over time for the three tested bacteria, S. clausii, the adapted S. clausii and S. patagoniensis. Fully grown bacterial suspensions were used at day 0. A starting population is maintained at least until day 3, but after 7 days in an anoxic environment the population rapidly reduces in size and a small stable population is maintained. Opening the bottles at day 7 leads to an increase in total cell counts for all strains when measured at 14 days (rows in bold).*

| **Date** | **Species** | **Gate** | **Average** | |
|---|---|---|---|---|
| 0 | *S. clausii* | Intact | 2.0E+05 | ± 1.1E+05 |
| 0 | *S. clausii* | Damaged | 9.1E+04 | ± 4.9E+04 |
| 0 | *S. clausii t10* | Intact | 5.4E+05 | ± 3.6E+04 |
| 0 | *S. clausii t10* | Damaged | 3.1E+05 | ± 4.0E+04 |

| | | | | |
|---|---|---|---|---|
| 0 | *S. patagoniensis* | Intact | 7.4E+05 | ± 1.7E+05 |
| 0 | *S. patagoniensis* | Damaged | 5.4E+05 | ± 1.2E+05 |
| 3 | *S. clausii* | Intact | 1.8E+05 | ± 4.5E+04 |
| 3 | *S. clausii* | Damaged | 5.6E+04 | ± 2.2E+04 |
| 3 | *S. clausii t10* | Intact | 6.8E+05 | ± 1.9E+04 |
| 3 | *S. clausii t10* | Damaged | 5.3E+04 | ± 2.1E+04 |
| 3 | *S. patagoniensis* | Intact | 7.5E+05 | ± 5.6E+04 |
| 3 | *S. patagoniensis* | Damaged | 7.4E+04 | ± 4.4E+04 |
| 7 | *S. clausii* | Intact | 1.8E+05 | ± 3.1E+04 |
| 7 | *S. clausii* | Damaged | 1.1E+05 | ± 2.4E+04 |
| 7 | *S. clausii t10* | Intact | 6.7E+05 | ± 5.3E+04 |
| 7 | *S. clausii t10* | Damaged | 1.1E+05 | ± 1.6E+04 |
| 7 | *S. patagoniensis* | Intact | 5.5E+05 | ± 2.0E+05 |
| 7 | *S. patagoniensis* | Damaged | 1.7E+05 | ± 6.3E+04 |
| 14 | *S. clausii* | Intact | 6.1E+04 | ± 2.7E+04 |
| 14 | *S. clausii* | Damaged | 1.7E+05 | ± 3.3E+04 |
| **14** | ***S. clausii*** | **Intact** | **1.5E+05** | ± **1.5E+03** |
| **14** | ***S. clausii*** | **Damaged** | **9.5E+04** | ± **3.6E+04** |
| 14 | *S. clausii t10* | Intact | 1.8E+05 | ± 3.4E+04 |
| 14 | *S. clausii t10* | Damaged | 1.3E+05 | ± 1.1E+04 |
| **14** | ***S. clausii t10*** | **Intact** | **2.6E+05** | ± **6.0E+04** |
| **14** | ***S. clausii t10*** | **Damaged** | **2.9E+05** | ± **3.1E+02** |
| 14 | *S. patagoniensis* | Intact | 2.3E+05 | ± 2.5E+04 |

| | | | | |
|---|---|---|---|---|
| 14 | *S. patagoniensis* | Damaged | 2.4E+05 | ± 7.4E+04 |
| **14** | ***S. patagoniensis*** | **Intact** | **1.7E+05** | ± **4.0E+04** |
| **14** | ***S. patagoniensis*** | **Damaged** | **3.9E+05** | ± **3.3E+04** |

*Table S 3 – Detailed calculations for the estimation of $CO_2$ and dissolved inorganic carbon in the system at day 14.*

| | **Sample** | **#** | **Opened at day 7** | **$CO_2$ concentration (%) m[CO2]h** | **Pressure in bottle (kPa)** | **Total pressure (kPa)** | **Total pressure (atm)** | **Partial pressure $CO_2$ (atm) $pCO_2$** | **$CO_2$ concentration in headspace normalized to liquid volume (mol/L)** |
|---|---|---|---|---|---|---|---|---|---|
| | | | | | | $P_T$ (kPa) = 101.325 + pressure in bottle | $P_T$ (atm) = $P_T$ / 101.3 | $pCO_2$ (g) = m[CO2]$_h$ /100 * $P_T$ | $[CO_2]_h$ (g) = n/V = partial pressure*($V_h$/$V_l$)/(R*T) |
| 1 | Cont | 1 | n | 0 | -23.0 | 78.325 | 0.77 | 0.0000 | 0.00E+00 |
| 2 | Cont | 2 | n | 0 | -27.8 | 73.525 | 0.73 | 0.0000 | 0.00E+00 |
| 3 | Cont | 3 | y | 0 | -13.6 | 87.725 | 0.87 | 0.0000 | 0.00E+00 |
| 4 | Cont | 4 | y | 0 | -18.4 | 82.925 | 0.82 | 0.0000 | 0.00E+00 |
| 5 | *Sc* | 1 | n | 0 | -22.4 | 78.925 | 0.78 | 0.0000 | 0.00E+00 |
| 6 | *Sc* | 2 | n | 0 | -31.7 | 69.625 | 0.69 | 0.0000 | 0.00E+00 |
| 7 | *Sc* | 3 | y | 0 | -19.4 | 81.925 | 0.81 | 0.0000 | 0.00E+00 |
| 8 | *Sc* | 4 | y | 0 | -6.3 | 95.025 | 0.94 | 0.0000 | 0.00E+00 |
| 9 | *Scg10* | 1 | n | 0 | -43.0 | 58.325 | 0.58 | 0.0000 | 0.00E+00 |
| 10 | *Scg10* | 2 | n | 0 | -43.1 | 58.225 | 0.57 | 0.0000 | 0.00E+00 |
| 11 | *Scg10* | 3 | y | 0 | -19.2 | 82.125 | 0.81 | 0.0000 | 0.00E+00 |
| 12 | *Scg10* | 4 | y | 0.0325 | -18.9 | 82.425 | 0.81 | 0.0003 | 2.14E-05 |
| 13 | *Sp* | 1 | n | 0.3884 | -34.4 | 66.925 | 0.66 | 0.0026 | 2.08E-04 |
| 14 | *Sp* | 2 | n | 0.2396 | -48.0 | 53.325 | 0.53 | 0.0013 | 1.02E-04 |
| 15 | *Sp* | 3 | y | 0.4783 | -11.5 | 89.825 | 0.89 | 0.0042 | 3.43E-04 |
| 16 | *Sp* | 4 | y | 1.0937 | -16.7 | 84.625 | 0.84 | 0.0091 | 7.39E-04 |

*Table S 3 – Detailed calculations for the estimation of $CO_2$ and dissolved inorganic carbon in the system at day 14 (continued).*

| | **Dissolved $CO_2$ (mol / Lt)** | **Concentration $H_2CO_3$ (mol / Lt)** | **pH** | **Concentration of $H^+$ from pH (mol / Lt)** | **Concentration $HCO_3^-$ (mol / Lt)** | **Concentration $CO_3^{2-}$ (mol / Lt)** | **Total dissolved inorganic carbon (mol / Lt)** | **Total inorganic carbon (mol / Lt)** |
|---|---|---|---|---|---|---|---|---|
| | $[CO_2]_{aq}$= $pCO_2$* 0.034 (mol/L.atm) | $[H_2CO_3]$ = $[CO_2]_{aq}$*1.7x10$^{-3}$ | | $[H^+]$ = $10^{-pH}$ | $[HCO_3^-]$ = ($H_2CO_3/H^+$) * 2.5x10$^{-4}$ | $[CO_3^{2-}]$ =($HCO_3^-$ /$H^+$)*4.7x10$^{-11}$ | = $H_2CO_3$ + $HCO_3^-$ + $CO_3^{2-}$ + $CO_2$ (*l*) | $[C_{inorganics}]$= $H_2CO_3$ + $HCO_3^-$ + $CO_3^{2-}$+ $CO_2$ (*l*) + $CO_2$ (*g*) |
| 1 | 0.00E+00 | 0.00E+00 | 10.94 | 1.14815E-11 | 0.00E+00 | 0.00E+00 | 0.00E+00 | 0.00E+00 |
| 2 | 0.00E+00 | 0.00E+00 | 10.92 | 1.20226E-11 | 0.00E+00 | 0.00E+00 | 0.00E+00 | 0.00E+00 |
| 3 | 0.00E+00 | 0.00E+00 | 10.90 | 1.25893E-11 | 0.00E+00 | 0.00E+00 | 0.00E+00 | 0.00E+00 |
| 4 | 0.00E+00 | 0.00E+00 | | 1 | 0.00E+00 | 0.00E+00 | 0.00E+00 | 0.00E+00 |
| 5 | 0.00E+00 | 0.00E+00 | 9.28 | 5.24807E-10 | 0.00E+00 | 0.00E+00 | 0.00E+00 | 0.00E+00 |
| 6 | 0.00E+00 | 0.00E+00 | 9.43 | 3.71535E-10 | 0.00E+00 | 0.00E+00 | 0.00E+00 | 0.00E+00 |
| 7 | 0.00E+00 | 0.00E+00 | 9.28 | 5.24807E-10 | 0.00E+00 | 0.00E+00 | 0.00E+00 | 0.00E+00 |
| 8 | 0.00E+00 | 0.00E+00 | 9.15 | 7.07946E-10 | 0.00E+00 | 0.00E+00 | 0.00E+00 | 0.00E+00 |
| 9 | 0.00E+00 | 0.00E+00 | 9.71 | 1.94984E-10 | 0.00E+00 | 0.00E+00 | 0.00E+00 | 0.00E+00 |
| 10 | 0.00E+00 | 0.00E+00 | 9.63 | 2.34423E-10 | 0.00E+00 | 0.00E+00 | 0.00E+00 | 0.00E+00 |
| 11 | 0.00E+00 | 0.00E+00 | 9.27 | 5.37032E-10 | 0.00E+00 | 0.00E+00 | 0.00E+00 | 0.00E+00 |
| 12 | 8.99E-06 | 1.53E-08 | 9.22 | 6.02560E-10 | 6.34E-03 | 4.94E-04 | 6.84E-03 | 6.854E-03 |
| 13 | 8.72E-05 | 1.48E-07 | 9.54 | 2.88403E-10 | 1.29E-01 | 2.09E-02 | 1.50E-01 | 1.496E-01 |
| 14 | 4.29E-05 | 7.29E-08 | 9.63 | 2.34423E-10 | 7.77E-02 | 1.56E-02 | 9.33E-02 | 9.339E-02 |
| 15 | 1.44E-04 | 2.45E-07 | 9.13 | 7.41310E-10 | 8.27E-02 | 5.23E-03 | 8.80E-02 | 8.821E-02 |
| 16 | 3.11E-04 | 5.28E-07 | 9.06 | 8.70964E-10 | 1.52E-01 | 8.16E-03 | 1.60E-01 | 1.604E-01 |

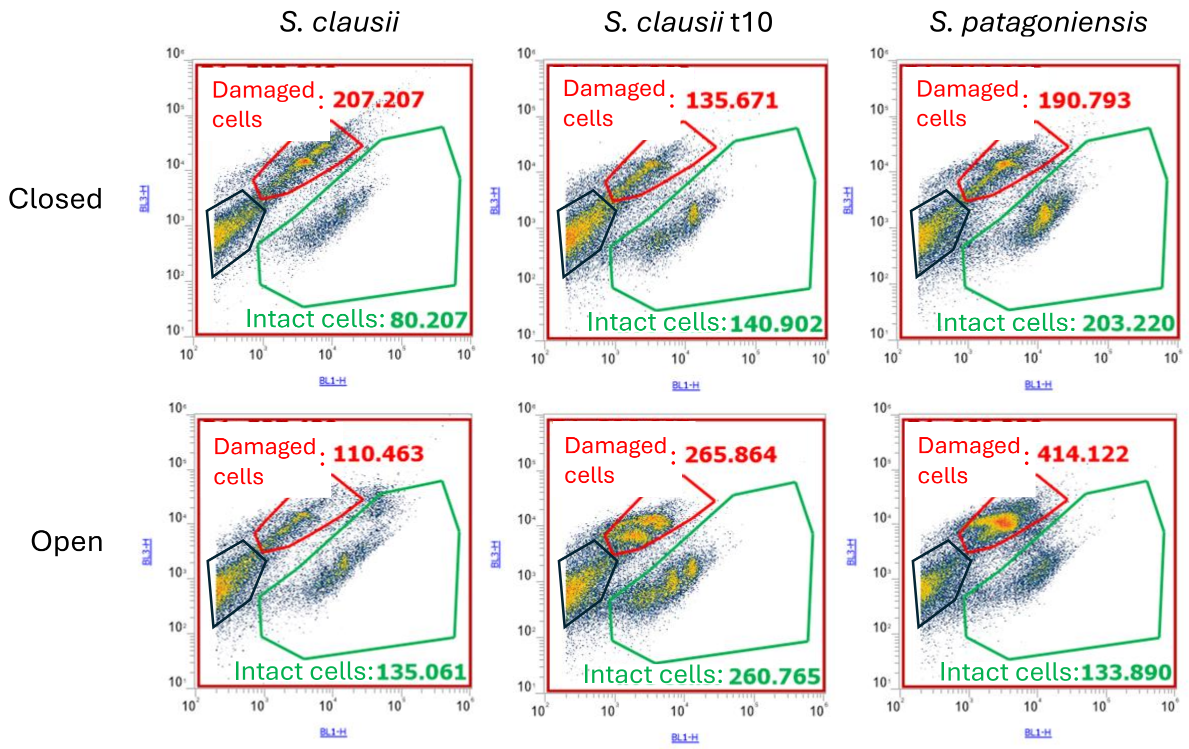


*Figure S 7 – Density plots of flow cytometry measurements for all three strains at day 14. Red gate: damaged cells; green gate: intact cells; blue gate: background. Top plots are from the bottles that remained closed while bottom plots are from the bottles that were opened at day 7. A noticeably denser cloud, for damaged and intact cells, is visible in the opened bottles as compared to the closed ones.*

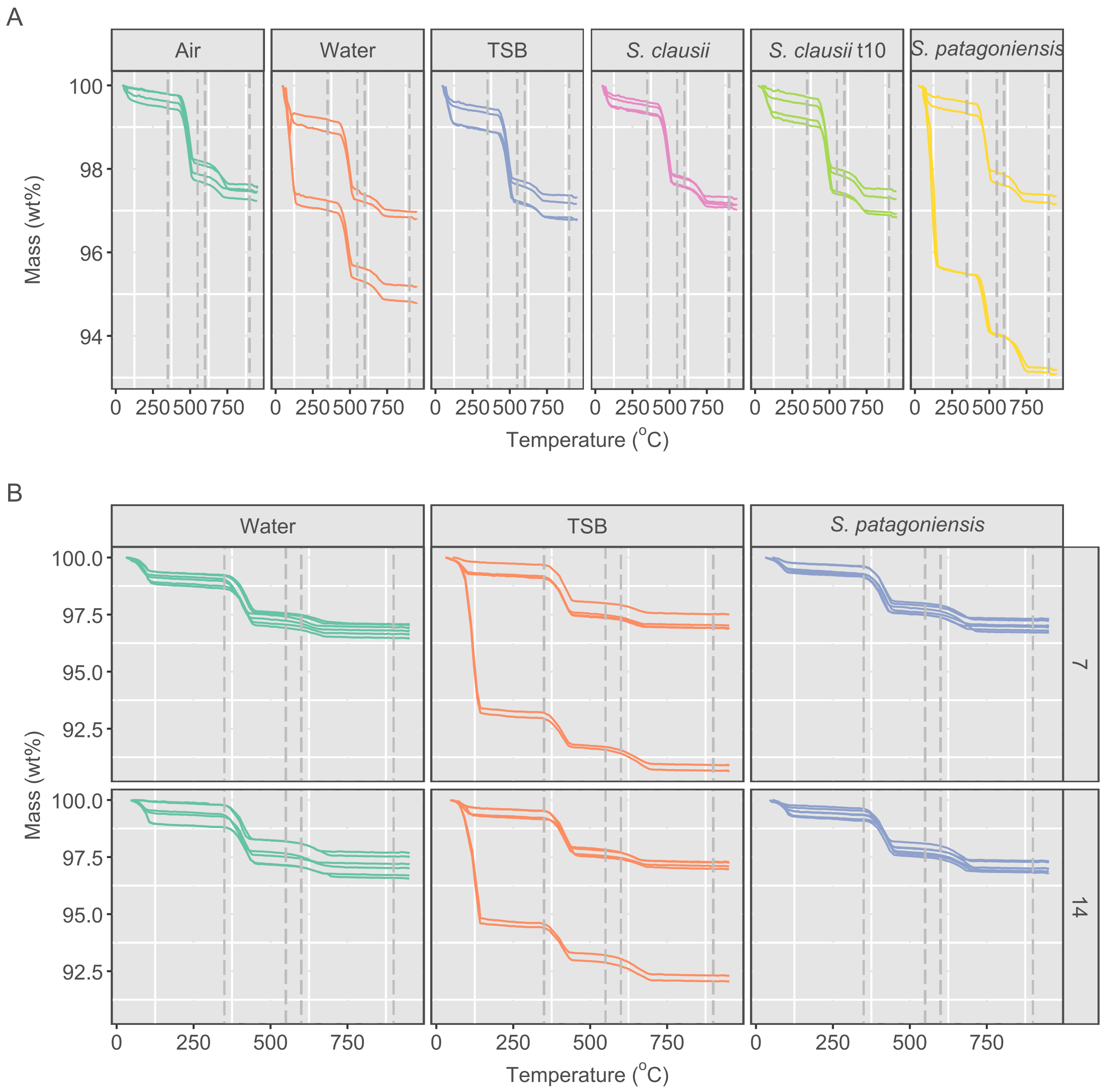


*Figure S 8 – A) Thermogravimetry analysis was performed on four different lime mortar samples at 7 days for six different applications. B) Thermogravimetry analysis was performed on six different lime mortar samples at 7 and 14 days for three different applications. Three stages of mass loss are observed: 20 – 100 °C which is the release of free $H_2O$, 350 – 550 °C which is the release of $H_2O$ from $Ca(OH)_2$ and 650 -950 °C for the release of $CO_2$ from $CaCO_3$.*

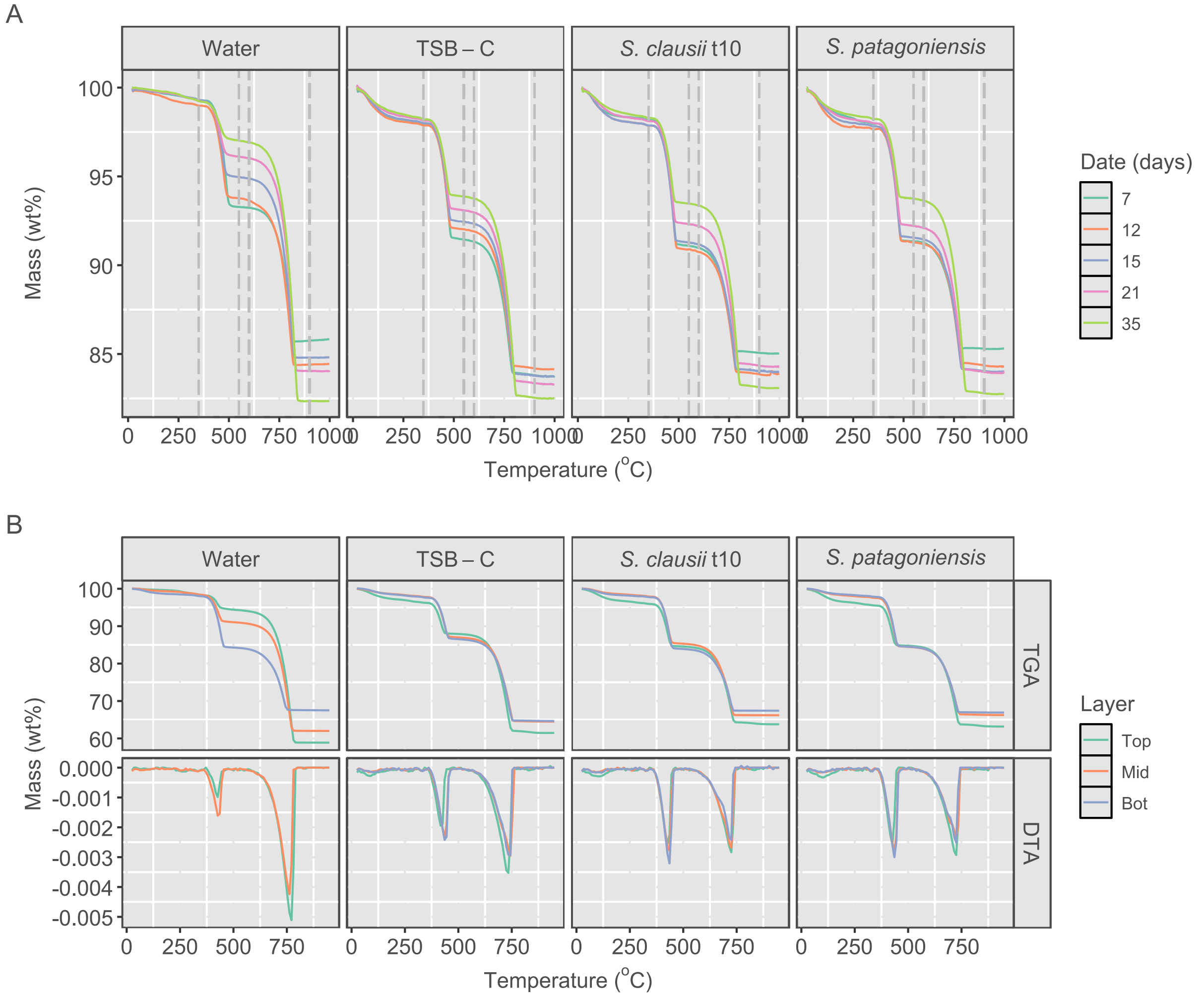


*Figure S 9 – Thermogravimetry analysis was performed on lime pastes at 7, 12, 15, 21 and 35 days for four different applications. B) Thermogravimetry analysis was performed on lime pastes samples at 21 days for four different applications. Each sample was grinded into three layers, a top, mid and bottom layer. Three stages of mass loss are observed: 20 – 250 °C which is the release of $H_2O$ and burning of organic matter, 350 – 550 °C which is the release of $H_2O$ from $Ca(OH)_2$ and 650 -950 °C for the release of $CO_2$ from $CaCO_3$.*

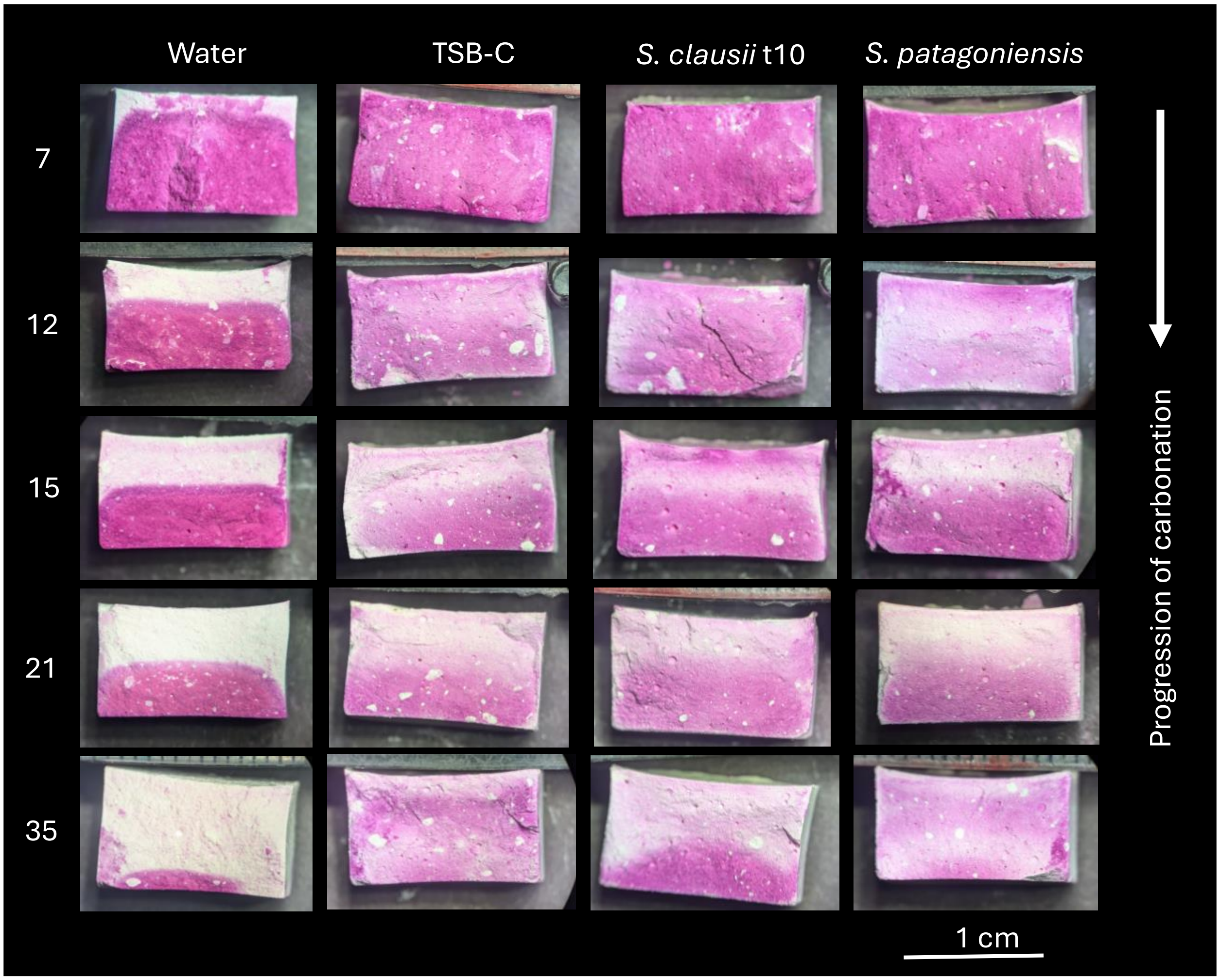


*Figure S 10 – Progression of carbonation as measured with spraying of phenolphthalein on a representative sample per condition and date. Arrow indicates direction of $CO_2$ diffusion, and all samples are 1.4 cm wide.*
